# GADMA: Genetic algorithm for inferring demographic history of multiple populations from allele frequency spectrum data

**DOI:** 10.1101/407734

**Authors:** Ekaterina Noskova, Vladimir Ulyantsev, Klaus-Peter Koepfli, Stephen J. O’Brien, Pavel Dobrynin

## Abstract

The demographic history of any population is imprinted in the genomes of the individuals that make up the population. One of the most popular and convenient representations of genetic information is the allele frequency spectrum or AFS, the distribution of allele frequencies in populations. The joint allele frequency spectrum is commonly used to reconstruct the demographic history of multiple populations and several methods based on diffusion approximation (e.g., *∂*a*∂*i) and ordinary differential equations (e.g., *moments*) have been developed and applied for demographic inference. These methods provide an opportunity to simulate AFS under a variety of researcher-specified demographic models and to estimate the best model and associated parameters using likelihood-based local optimizations. However, there are no known algorithms to perform global searches of demographic models with a given AFS. Here, we introduce a new method that implements a global search using a genetic algorithm for the automatic and unsupervised inference of demographic history from joint allele frequency spectrum data. Our method is implemented in the software GADMA (Genetic Algorithm for Demographic Analysis, https://github.com/ctlab/GADMA). We demonstrate the performance of GADMA by applying it to sequence data from humans and non-model organisms and show that it is able to automatically infer a demographic model close to or even better than the one that was previously obtained manually. Moreover, GADMA is able to infer demographic models at different local optima close to the global one, making it is possible to detect more biology corrected model during further research.

## Introduction

To understand the evolution of species and their populations, it is important to understand what events occurred in their past and when. The genetic diversity and structure of species are shaped by the combined processes of changes in effective population size, population divergence and/or migration (gene flow) operating over the course of thousands or millions of years. Records of population history are imprinted in the genomes of individuals within species and this history can be inferred using a a variety of algorithmic and statistical methods. With the rise of next-generation sequencing (NGS) technologies and abundant genome data, it has become possible to explore complex and parameter-rich demographic models that include the estimation of mutation rate, changes in population size, nonrandom mating, admixture, and selection (Pool *et al.*, 2010; Schraiber and Akey, 2015). However, given the infinitely large number of permutations at which these processes operate over various time intervals, there is no method that can guarantee to find the best demographic model that best fits the observed data.

One of the primary methods for inferring demographic models from genomic data is based on the allele frequency spectrum (AFS), also known as the site frequency spectrum (Beichman *et al.*, 2018; Schraiber and Akey, 2015). In essence, the AFS describes the distribution of derived allele frequencies of biallelic loci (SNVs) in a population or sample of populations based on the number of sequenced chromosomes (Fisher, 1931). An AFS can provide information about how the populations developed based on observed genetic variation sampled from current individuals of those populations. Many studies have been devoted to testing and understanding the behavior of allele-frequency spectra under different demographic scenarios (Adams and Hudson, 2004; Chen *et al.*, 2007; Marth *et al.*, 2004; Myers *et al.*, 2008; Voight *et al.*, 2005).

Two of the most popular methods of historical demographic inference based on AFS are the faster continuous-time sequential Markovian coalescent approximation (fastsimcoal2, (Excoffier and Foll, 2011)) and the diffusion approximation (*∂*a*∂*i, (Gutenkunst *et al.*, 2009)). fastsimcoal2 can handle any number of populations, but it is computationally hard because it simulates multiple AFS simultaneously to estimate the most stable one. *∂*a*∂*i simulates AFS using a numerical solution of the partial diffusion equation (PDE), which corresponds to the presented demographic model and then provides the likelihood of the model (Figure 1). Unfortunately, PDE leads to some computational difficulties associated with analyses of complicated demographic models and large sample sizes. As a result, *∂*a*∂*i can only handle up to three populations. More recently, two new methods called *moments* (Jouganous *et al.*, 2017) and momi2 (Kamm *et al.*, 2018) has been introduced. *moments* like *∂*a*∂*i is based on two models of population genetics: the Wright-Fisher generation model and the infinite sites mutational model. momi2 is based on another generation model — the Moran model. *moments* uses ordinary differential equations to simulate AFS, which is faster and more stable than diffusion approximation in *∂*a*∂*i, based on simulations comparing the two methods (Jouganous *et al.*, 2017). *moments* presents a tradeoff between speed and accuracy in AFS-based demographic inference, can handle up to five populations and provides the same interface as *∂*a*∂*i.

**FIG. 1.**
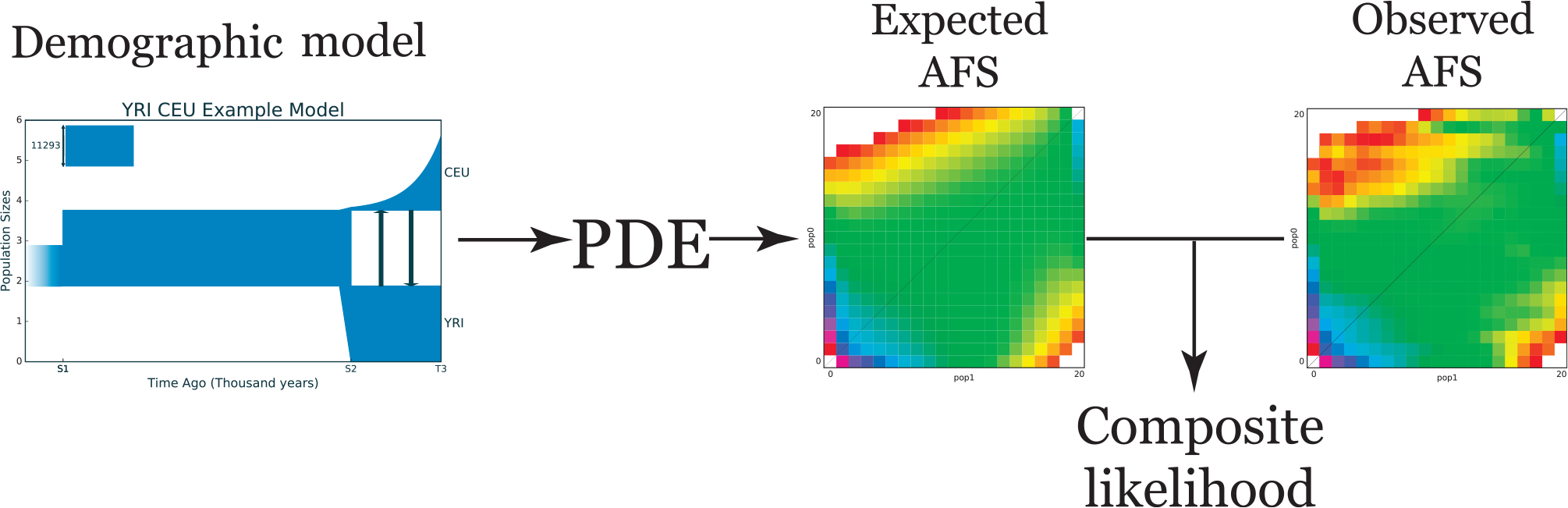
General scheme of *∂*a*∂*i. In order to compare a demographic model and the real observed AFS, *∂*a*∂*i extracts the expected AFS from the demographic model using a numerical solution of the partial differential equation (PDE) that corresponds to that demographic model, and calculates the composite likelihood between the expected and observed allele frequency spectrum.

Ideally, researchers seek to find the model of demographic history that best describes or “fits” their data. *∂*a*∂*i and *moments* provide an opportunity to run multiple optimizations to help fit parameters of a given demographic model that maximizes the value of the composite likelihood. But optimizations based on gradient descent, for example, the Broyden-Fletcher-Goldfarb-Shanno (BFGS) algorithm (Broyden, 1970; Fletcher, 1970; Goldfarb, 1970; Shanno, 1970) or its modifications, use numerical differentiation and are ineffective in practice, because of the complex structure of the search space for demographic models. Optimizations may also be inefficient due to another set of methods that offer existing solutions based on local search algorithms without gradients, such as the Nedler and Mead method (Nelder and Mead, 1965) or Powells method (Powell, 1964). As a result, all existing optimizations find local optima close to the initial values and require many runs to be performed using different initial model parameters, most of which are unknown or lack empirical data. Despite these drawbacks, *∂*a*∂*i and *moments* are efficient instruments, since they can simulate an allele frequency spectrum from any demographic model. In other words, the problem of finding a demographic model from the allele frequency spectrum is the inverse problem, which can be solved by solving the direct problem, that is simulation of the AFS from a given demographic model, with approximate numerical methods, such as diffusion or *moments* approximations. The lack of accurate and rapid differentiation and the complex structure of the search space led us to consider the usage of global optimization methods, such as the genetic algorithm.

The genetic algorithm (GA) (Holland, 1975) is one of the most efficient heuristic algorithms for global searching of complex and rich parameter space. Its primary application is optimization of a fitness function, which is either not differentiable or it cannot be differentiated in a sufficiently effective way, for example, when the function is not representable in “closed-form expression”. GA is based on the principle of evolution and simulates natural selection using operations of “mutation” and “crossover”, which ideally results in the most adapted individual, the one which has the best value of the fitness function. The versatility of the GA has led to its wide application, including reconstruction of phylogenetic trees (Zwickl, 2006), ancestral genome composition inference (Gao *et al.*, 2013), and evolutionary biology in general (Hamblin, 2013).

In this paper we present a new method based on the genetic algorithm to automatically infer the best fitting demographic model from AFS data for two or three populations. Our method assumes the ability to simulate the AFS from the demographic model, for example, using either *∂*a*∂*i or *moments*. The GA framework overcomes limitations of local search optimizations and is more flexible in handling the complexity of demographic models by allowing an increase in the number of parameters and estimation of parameters such as functions population size changes that are sudden, linear or exponential. We implemented our method in the software GADMA (Genetic Algorithm for Demographic Model Analysis), which is written in Python and available at Github (https://github.com/ctlab/GADMA).

## Materials and Methods

This section provides definitions of the allele frequency spectrum and the composite likelihood scheme that are used in existing optimizations (*∂*a*∂*i and *moments*) and implemented in our method. After this background, the problem of demographic model search from observed AFS data is formulated in terms of computer science. We then introduce a developed representation of a demographic model for a general approach to our method, including the genetic algorithm with its operations of “mutation” and “crossover”.

### Basic definitions and concepts

Assume there are *P* populations and for each population *i* there exists information about *n*_*i*_ chromosomes. The AFS is the *P*-dimensional array *A*, where each entry *A*[*d*_1_,…,*d*_P_] ∈ ℕ, *d*_*i*_ ∈ [0,*n*_*i*_], ∀*i* ∈ [1,*P*] records the number of SNVs (relative to the common reference genome), which are exactly seen at *d*_1_ chromosomes from population 1, *d*_2_ chromosomes from population 2, … and *d*_*P*_ chromosomes from population *P*. For example, if we have two populations, then the AFS is a two-dimensional matrix, where *A*[*i*,*j*] represents the number of polymorphisms that occurred exactly in *i* individuals of the first population and in *j* individuals in the second population (Figure 2). Several examples of allele frequency spectra are presented in Figure 3.

**FIG. 2.**
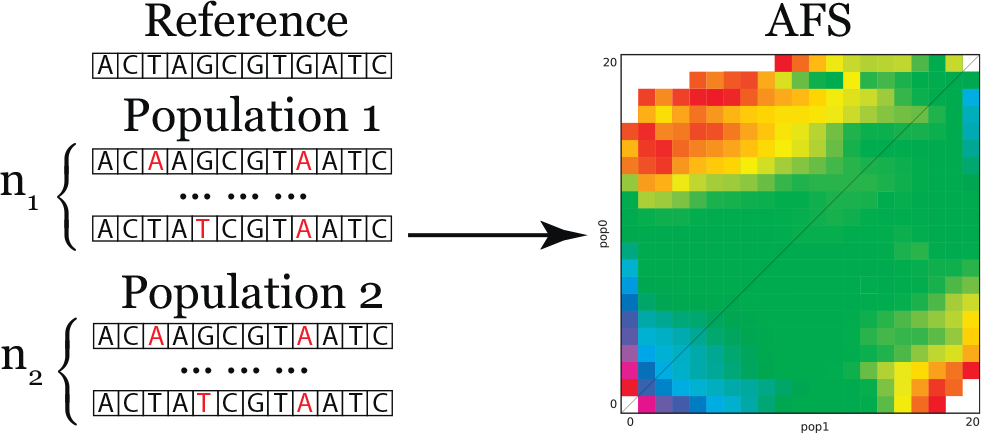
Diagram showing the process of building the allele frequency spectrum. *P* populations are observed (*P* = 2 in the figure), then for each population *i* there is genetic information for *n*_*i*_ chromosomes. To build the allele frequency spectrum we need to call SNVs (relative to some reference) and calculate their frequencies.

**FIG. 3.**
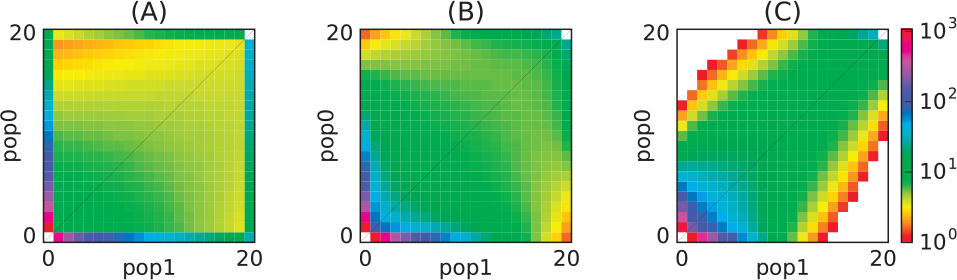
Examples of different allele frequency spectra. (A)Isolation of a population after divergence; (B) Low symmetrical migrations between populations following divergence; (C) High asymmetrical rates of migrations between populations following divergence.

Assume that we can simulate the AFS *M* from the proposed demographic model. Assuming no linkage between derived alleles, each element of the allele frequency spectrum *S*[*d*_1_,…,*d*_*P*_] is an independent Poisson value with an mean equal to *M* [*d*_1_,…,*d*_*P*_]. We then calculate the likelihood – the probability of obtaining the observed *S*, if the expectation spectrum is *M*, as the product of (*n*_1_ +1)(*n*_2_ +1)…(*n*_*P*_ +1) Poisson likelihood functions:

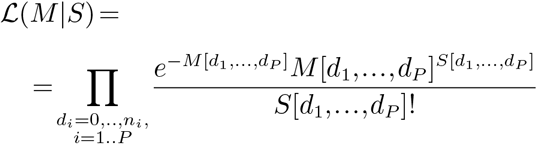

In the case of linked alleles, 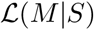 is the composite likelihood. Demographic models inferred by *∂*a*∂*i and *moments* can be compared by computing the 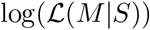. Because 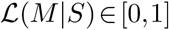, then 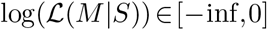 and the greater is the log-likelihood, the better the model fits the observed AFS. In this paper, 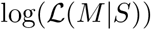 was chosen as the fitness function of the genetic algorithm, as discussed below.

### Formulation of the problem

Consider a function *f* (Θ,*A*,*C*) that takes the parameters 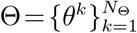, *k* ∈ ℝ, the AFS *A* ∈ ℝ^*P*×*P*^, the set of constants 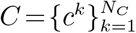 and returns the measure of the correspondence between the parameters Θ and the AFS *A*.

The function *f* (Θ,*A*,*C*) builds a demographic model with respect to the parameters Θ that unambiguously determine it, calculates the expected AFS *M* with respect to the constants *C*, and then determines the degree of similarity between *M* and the observed *A* by the composite likelihood. The constants can be various parameters of algorithms for calculating the expected AFS, such as grid sizes for the numerical solution of a differential equation in *∂*a*∂*i, or population models parameters, including the average number of new mutated sites per individual in a generation *θ*_0_ for the infinite-sites mutational model, or the time *T*_*g*_ for one generation in the Wright-Fisher model. The function *f* can have different implementation details. Here *∂*a*∂*i and *moments* were selected for this purpose.

The purpose of this work is to develop an algorithm to search for the demographic model that best corresponds to the observed AFS. Formally, the problem can be formulated as follows:

**Input**

- *A* ∈ ℝ^*P*×*P*^ — the *P*-dimensional array, *P* ∈ {2,3}.
- 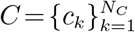 — the set of constants.

**Output**

- The set 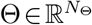 of values, that maximize the value of *f*:

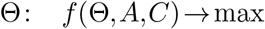

There are approximate solutions of this problem with an additional input — a demographic model with a fixed number of parameters, using various local search algorithms, but in practice, as mentioned above, these algorithms have proven to be ineffective. We present a new algorithm for the approximate solution of this more general problem using one of the most effective methods of global optimization — the genetic algorithm.

### Representation of the demographic model and its structure

Assume a division of one population into two new isolated subpopulations. Then the number of population splitting events directly depends on *P*, the number of considered populations. We represent the demographic model as a sequence of “time intervals” and population splits, each of which has a fixed number of parameters. Assume a fixed temporal order of the current observed populations: from an ancient population to the more recently formed population. This temporal order is usually known or can be imputed. Then if the number of populations is ≤3, then each split will divide the last formed population. Thus, a splitting event has only one parameter — the fraction of the population, which separates to form a new population.

The next important component of the demographic model is the concept of “time interval”. First, we define this as a segment of time during which a certain dynamic of change of effective size is maintained for each population. We consider three main demographic dynamics of population growth: sudden, linear and exponential change of the effective population size (Figure 4). Sudden change is very popular for its simplicity, but exponential change is a commonly used model for population growth as well. We include linear change as it is tradeoff between sudden and exponential change and is more realistic than sudden change. Secondly, the parameters of migration rates between populations are constant during a given time interval. Thus, each time interval has the following parameters:

- time,
- effective population sizes at the end of the time interval,
- demographic dynamics of effective population size change,
- migration between populations, if there is more than one population.

**FIG. 4.**
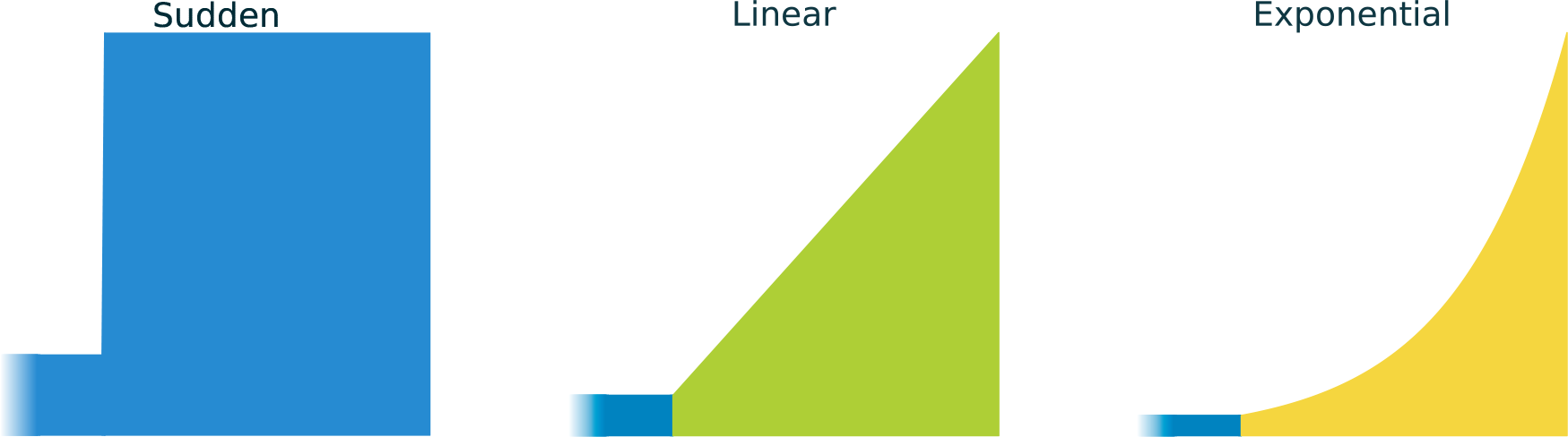
Diagrams of three primary demographic dynamics of population size change. Sudden population growth is very popular for applications in demographic models, as well as exponential population growth. Linear population growth is a tradeo between sudden size change and exponential change and it is also more realistic that sudden population growth. Different colors are used to highlight different types of population growth.

The sizes of the populations at the beginning of any time interval is equal to the number of populations at the end of the previous interval. The first time interval is a special one: we consider that it lasts from the beginning of the existence of the species and assume a demographic dynamic of sudden change for the effective population size of the ancestral population (Gutenkunst *et al.*, 2009). Therefore, in this interval the only parameter is the size of the ancestral population.

Note that the number of splitting events is determined by the number of populations under consideration, but the number of intervals can be varied and thus change the number of parameters of the demographic model, its detail, and complexity. We now can define the concept of the structure of the demographic model:

#### Definition 1

Structure of the demographic model — the number of time intervals that occur before and after a single splitting event, in the case of two populations. In the case of three populations, the number of time intervals prior to the first split, those between the first and second split, and the ones after the second split. For example, assume we observe three populations. At the beginning, there was an ancestral population (*P*_*A*_) and this population started to grow in effective size. Then a split occurred that divided this ancestral population into two daughter populations (*P*_1_ and *P*_2_) that changed in effective size during one interval, followed by a split of the second population (into *P*_2*a*_ and *P*_2*b*_), resulting in three descendant populations that changed within one subsequent time interval. The structure of such a model would be described as (2,1,1) (Figure 5). The simplest model structures would be for two populations – (1,1), and for three populations – (1,1,1).

**FIG. 5.**
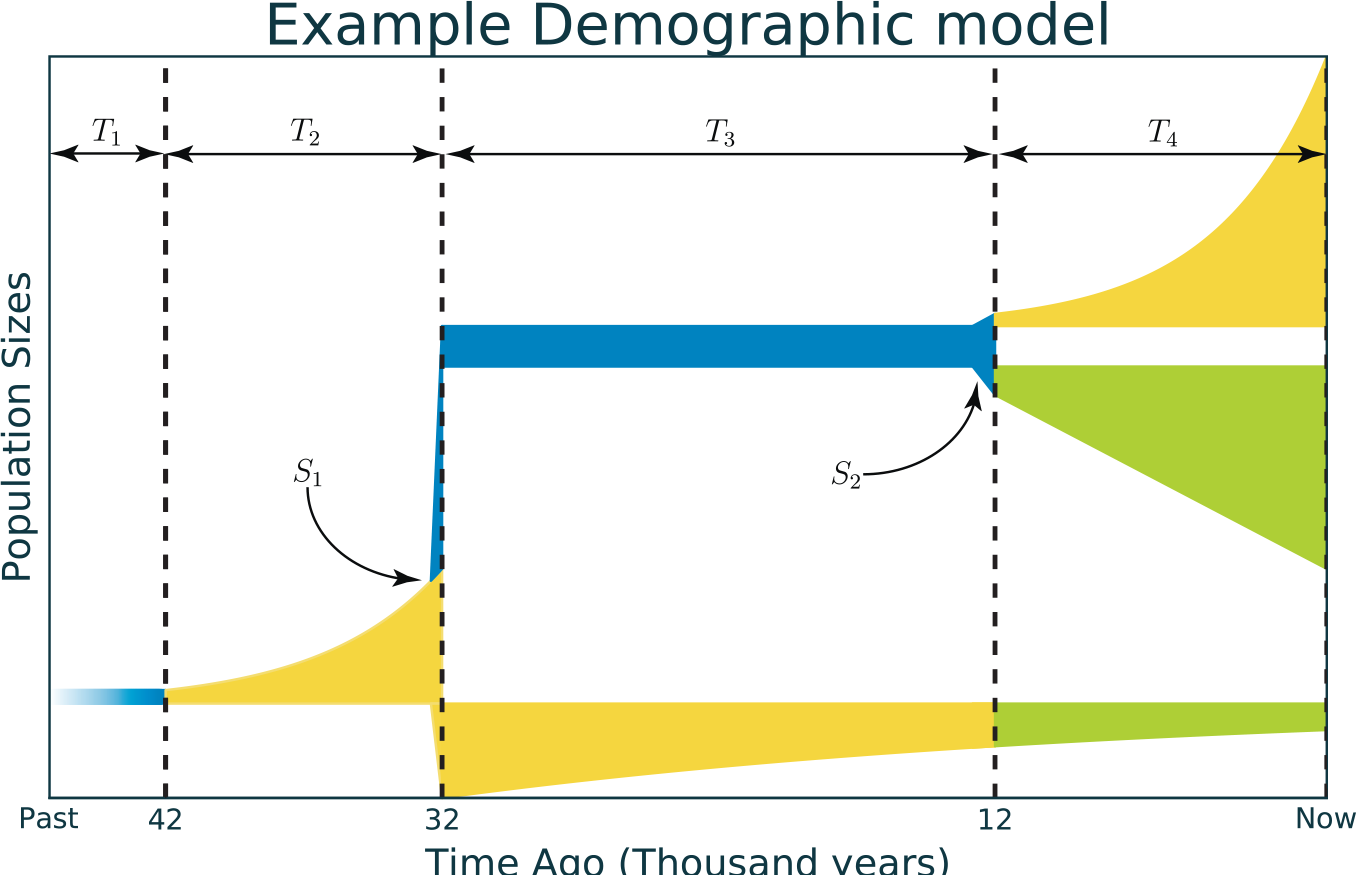
Example of representation of a demographic model with a (2,1,1) structure. Time is shown along the x-axis and population size on the y-axis. Four time intervals are shown: *T*_1_, *T*_2_, *T*_3_ and *T*_4_ and two populations splits: *S*_1_ and *S*_2_. The structure of this model is (2,1,1) because *T*_1_ and *T*_2_ are time intervals for one ancestral population, followed by split *S*_1_, *T*_3_ is the time interval for two populations, and *T*_4_ is the time interval for three final populations after the second split *S*_2_.

More formally, the structure of the model is a sequence of the form 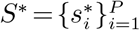, 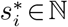 where *P* ∈{2,3} is the number of populations. In this case, the number of parameters *N*_Θ_(*S*\*) of the demographic model with the structure *S*\* will be determined as follows:

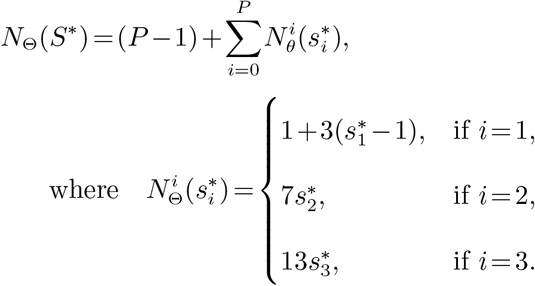

The term (*P* − 1) corresponds to the number of of split parameters, and 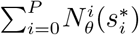 is the number of time intervals parameters. Thus, we can unambiguously interpret the demographic model according to the list of parameters and its structure by fixing for each time intervals a certain order of parameters.

### General approach

The general algorithm is a series of executions of the genetic algorithm (Figure 6). Suppose we have the initial and final structures that define the initial and final number of parameters (definition of the structure is presented in previous section) of the demographic models, derived from considerations about the populations we are trying to model. During the genetic algorithm, the structure of the model does not change; that is, the models parameters for the current structure are optimized. This restriction makes the procedure of crossover in the genetic algorithm simpler. As soon as the genetic algorithm stops, the parameters of the best-fitting model are adjusted in order to improve the likelihood with a local search algorithm. Then if the structure of the model is not quite complex, its complexity (in terms of parameters) is increased and the genetic algorithm is run again for the new structure. The genetic algorithm and local search are executed until the best-fitting parameters of the model with final structure is obtained.

**FIG. 6.**
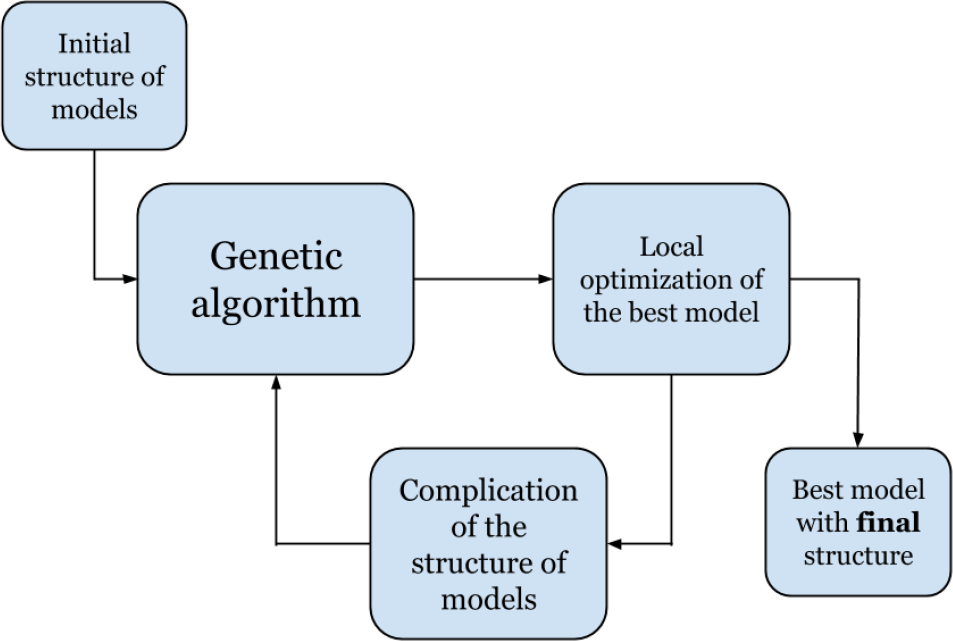
Diagram showing general algorithm used in GADMA and the relationship between the genetic algorithm that performs the global optimization search followed by the local optimization search.

#### Akaike information criterion

With an increase in the number of model parameters, we risk overfitting the model. A model with a large number of parameters will be better able to find parameter values corresponding to the observed data than a model with a smaller number of parameters, but at the same time it will correspond less to reality, for example, due to data errors. The Akaike information criterion (*AIC*) (Akaike, 1974) is commonly used to compare models with different numbers of parameters:

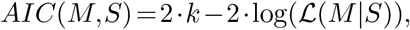

where *k* is number of parameters of the model and 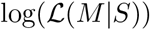 — if the value of the log-likelihood function. The smaller *AIC* is, the better the model fits the observed data.

Obviously, it is enough to compare only the final models for each structure after local search, since the number of parameters between the increases in model structure does not change, and therefore the value of *AIC* score depend only on the likelihood values. In the implementation of the GADMA method if the models with the best likelihood and best value of *AIC* score do not match, the user is informed about the overfitting.

### Genetic algorithm

The genetic algorithm is one of the most effective heuristic algorithms (Holland, 1975). It is based on the principles of evolution, where the aim of the algorithm is to find an approximate solution to a problem that has the maximum or minimum value of the fitness function. At the beginning of the algorithm, a fixed-sized set of random solutions, called individuals, is created. The set itself is called a generation. Each individual is assigned a value of fitness, which is determined by the value of the fitness function. After this, new generations are iteratively produced with the help of mutations, crossover and selection of the fittest individuals. All these operations can be either deterministic or random, and their order can vary in different implementations. In our case, individuals are demographic models of the same structure, and the fitness function is the log-likelihood 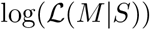, as described above.

In the first step of GADMA, a set of demographic models are randomly generated, if they have not been already specified. To form a new generation of demographic models, we select the most adapted models among a set of mutated, crossed, and random models in the previous generation. The value of the fitness function is used to select the most adapted models. The choice of models to be mutated or have crossover is random, but the probability of choice is directly proportional to the value of fitness: the better the fitness of the model is, the more likely it is to be selected. The genetic algorithm stops when it can no longer obtain a better demographic model by the value of the fitness function for several iterations (Figure 7).

**FIG. 7.**
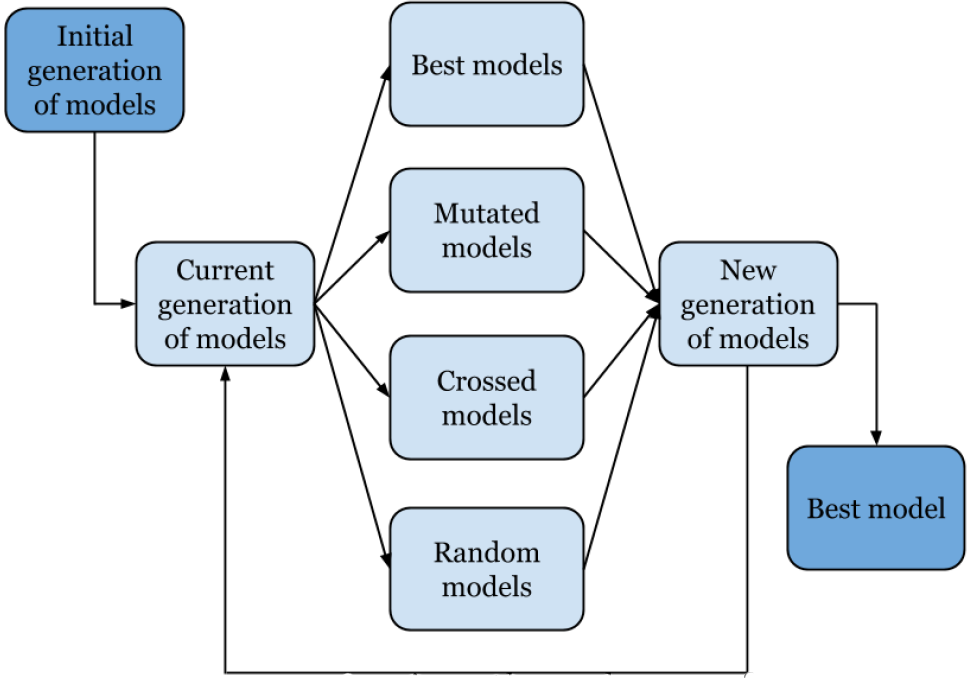
Diagram showing the scheme of the genetic algorithm used in GADMA.

#### Mutation of the demographic model

The mutation of the demographic model (Figure 8) is equal to the process of changing the values of several parameters. There are two constants associated with mutation of the model: the rate and the strength of mutation. The number of parameters to be mutated is sampled from a binomial distribution with a mean equal to the mutation strength. Parameters that are mutated are chosen with the probability that is directly proportional to their weights, which at the beginning are equal (that is, the choice is equally probable), and then each weight can be increased when a mutation of the corresponding parameter has occurred, which leads to an improvement in the model. The measure of how much the value of each parameter is mutated is determined by the sign (+1 or −1 equally probable) and the rate of the mutation that is randomly sampled from the normal distribution, with the mean equal to the mutation rate and a variance equal to half of the mean.

### Adaptive mutation rate and mutation strength

In the initial iterations of the genetic algorithm, strong mutations of a large number of parameters are much more effective than smaller mutations of a small number of parameters, whereas when approaching the optimal solution the opposite is true. Therefore, the rate and strength of the mutation can be adaptive, that is, it changes during the operation of the algorithm. There are several ways to make an adaptive value, one of the most popular being the one-fifth algorithm (Schumer and Steiglitz, 1968). First, we apply it to the mutation rate: at each iteration, if we have a successful solution, that is, it becomes better after mutation, then we multiply the mutation rate by the constant *C* ∈ [1,2]. If the solution is not “successful, then we divide by a fourth-degree root of *C*, decreasing the mutation rate. In the case of the mutation strength for the success of the solution, it is necessary to additionally check whether the decision has become the best solution during the entire course of the algorithms run.

Thus, often getting a new best solution with a mutation that occurs at the beginning of the genetic algorithm, we will increase the number of parameters that are changed during the mutation operation and the degree to which they are changed. As we approach the optimum solution and decrease in frequency, the number of parameters will also decrease and lead to a more accurate search. An increase in the number of parameters being mutated leads to a more efficient crossover. At the same time, the mutation rate is changed more frequently than the mutation strength, which makes the mutation procedure more effective.

**FIG. 8.**
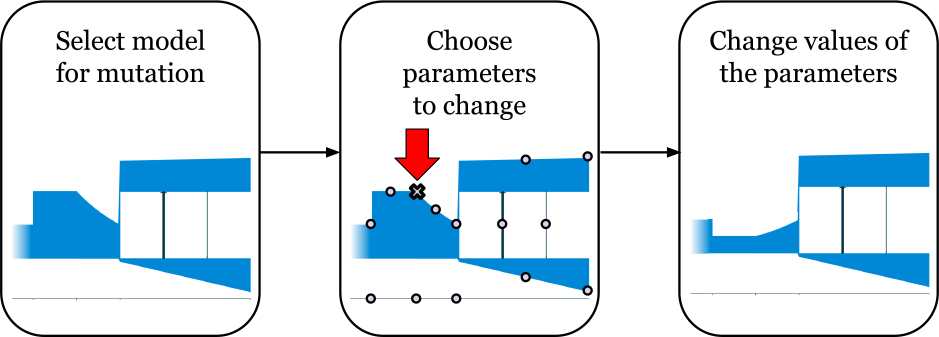
Diagram showing how mutation is used to change parameters of the demographic model in GADMA. In this example, only one parameter is mutated — ancestral population size at second time interval.

**FIG. 9.**
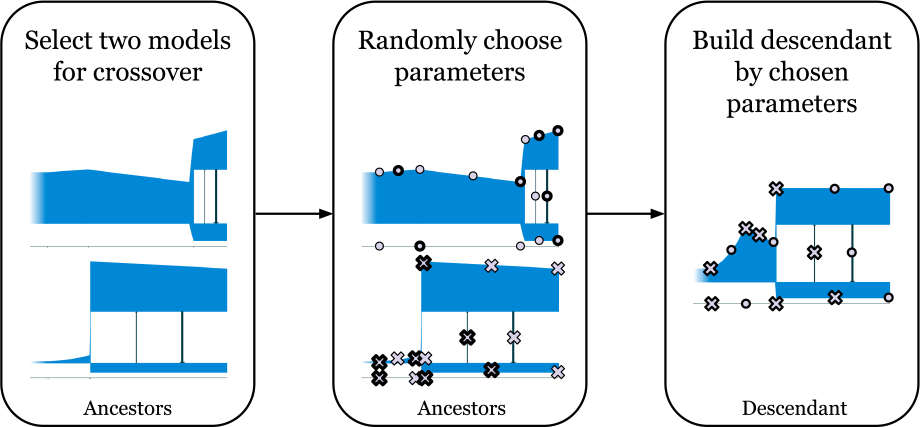
Diagram showing how crossover of two demographic models is performed in GADMA.

### Crossover of two demographic models

In order to have crossover of two demographic models, the models are represented as sequences of parameters. Each parameter of their descendant is chosen randomly with equal probability from one or the second parent (Figure 9). Since the structure of models do not change during the operation of the genetic algorithm, the number of parameters for all models will be the same. Consequently, the parameters can be unambiguously interpreted and easily crossed according to these criteria.

### Local search algorithms

Local optimizations are effective in cases when the initial solution is close to optimal. They are more accurate in adjusting parameters than the genetic algorithm, and can significantly improve its result. *∂*a*∂*i and *moments* provide the following choice of algorithms for local optimization:

- The Broyden-Fletcher-Goldfarb-Shanno (BFGS) algorithm.
- L-BFGS-B — a modification of the BFGS algorithm, which is more efficient when the optimal parameters are close to the bounds.
- The Nelder-Mead method or simplex-method.
- Powell’s method.

The first two methods are gradient descent optimization algorithms (Ruder, 2017) and the last two do not use a gradient. Powells method was proposed by the authors of *moments* and was noted as the most effective, so it was adapted for usage with *∂*a*∂*i and for our experimental studies it was chosen as a local search algorithm.

### Increasing the complexity of the structure of the demographic models

We need to be able to increase the complexity of the structure of the demographic model in order to find an optimal solution. To do this, a time interval is selected and then divided into two equal intervals (i.e., dividing based on the time median). The time interval is chosen randomly on the basis that the new structure *S\** should not become greater than the final *S*^*F*^ according to one of its values: 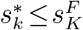, ∀*k* ∈ [1,*P*]. The values of the parameters of the new formed time intervals are calculated for the parent: the size of the population of the first time interval is equal to the size in the median of the parental time interval, and the parameters of the second time interval are equal to the population size at the end, with the time of both intervals equal to half of the parental time, while migration between populations and the demographic dynamics of population size change remain the same. In essence, the demographic model has more parameters after its structure is increased, however the history and likelihood remain the same.

## Results

We implemented the method described above in the program GADMA (Genetic Algorithm for Demographic Model Analysis), written in the Python programming language. We explored the efficiency of our method using several previously published data sets. GADMA was used to first infer demographic models for two and three modern human populations using the data set analyzed by (Gutenkunst *et al.*, 2009). For these analyses, different parameter values within GADMA were examined, including the initial structure of the demographic models and using either *∂*a*∂*i or *moments* to infer the optimal model of demographic history. We then inferred demographic models for the history of two populations of the Gillettes checkerspot butterfly, *Euphydryas gillettii*, and these were compared to the previously models reported by (McCoy *et al.*, 2014). Lastly, we used GADMA to reconstruct the demographic history of the Gaboon forest frog, *Scotobleps gabonicus*, which occurs in Central and Western African rainforests, based on a data set generated and originally reported by (Portik *et al.*, 2017).

### Testing the human Out of Africa model with GADMA

One of the most popular demographic history models for human populations is the so-called “Out of Africa” model, which consists of three populations (Gravel *et al.*, 2011; Gutenkunst *et al.*, 2009; Harris and Nielsen, 2013; Jouganous *et al.*, 2017):

- YRI — Yoruba individuals from Ibadan,
- CEU — Utah residents with ancestry from northern and western Europe,
- CHB — Han Chinese individuals from Beijing.

In order to demonstrate the effectiveness of our method, we choose to use the allele frequency spectrum from the paper of (Gutenkunst *et al.*, 2009), in which the *∂*a*∂*i method is introduced and the demographic history models for two (YRI, CEU) and three (YRI, CEU, CHB) populations are inferred from this spectrum. These models (Figure 10a and Figure 11a) were obtained from a large number of local optimization launches and also have a number of restrictions on the number of population parameters: the size of the YRI population does not change after the first expansion of the ancestral population, migrations are symmetrical and the dynamics of population size change are fixed as sudden, except for the last time interval for CEU and CHB, where exponential growth occurs. The model for two populations has a total of 6 parameters (Table 2) whereas that for three populations has a total of 14 (Table 4).

The 21 × 21 × 21 AFS was constructed in Gutenkunst et al. 2009 on the basis of the Environmental Genome Project (Sharp and Barrett, 2000). All biallelic SNVs from non-coding regions of 219 genes (totaling 5.01 Mb) were considered and the effective length of the used sequence was equal to *L* = 4.04 × 10^6^. We used the same neutral mutation rate equal to *μ* = 2.35 × 10^−8^ and the same generation time equal to *t*_*g*_ = 25 years as in Gutenkunst et al. 2009. Thus, the average frequency of mutation in one individual per generation is equal to *θ*_0_ = 4*μL* = 0.37976.

The following parameters of the genetic algorithm were used: the size of the generation of the demographic models was chosen to equal to 10, the strength and mutation rate was equal to 0.2, and the proportions of the best, mutated, crossed and random models in the new generation were 0.2 : 0.3 : 0.3 : 0.2. The strength and rate of mutation were adaptive with the constants of 1.05 and 1.02, respectively. The AFS was simulated using *∂*a*∂*i with a *G* = {40,50,60} grid size, the value of the likelihood was considered significant to the second decimal point, and the genetic algorithm stopped after 100 iterations without improvement. As for the local optimization search, the Powell method was chosen.

#### The YRI, CEU two population example

Three demographic models were inferred from the same AFS: using the same parameters as in (Gutenkunst *et al.*, 2009) (model 1) and two with all possible twelve parameters (models 2, 3) but with different initial demographic model structures. Model 2 had a structure (1,1), which then expanded to (2,1) during the GADMA run, whereas model 3 had an initial structure of (2,1). We ran GADMA 10 times for each of the three models (Table 2, Figure 10). All three models resulted in likelihoods better than the final demographic model inferred in Gutenkunst et al. 2009. The parameters of models 1 and 2 are not significantly different, and model 3 has the best likelihood value. Model 3 shows a lower population size of Europeans, a larger rate in their growth (from 25 individuals to 9 thousand) and a shorter separation time than the best model found in Gutenkunst et al. 2009 as well as models 1 and 2. Migration rates between the populations were chosen to be asymmetric, but they are largely equal to each other and coincide with values among the three models.

To demonstrate the inefficiency of the methods of local optimization for the model from (Gutenkunst *et al.*, 2009), one of the methods proposed by *∂*a*∂*i, BFGS, was launched 100 times. In each run, the initial parameters values were chosen randomly. The best value of the log-likelihood was *−*1629.24, which is quite far from the optimal value of *−*1066.35 reported in Gutenkunst et al. 2009. The average time of one optimization run was about 21 minutes. We then used a more efficient local optimization method implemented in the *dadi pipeline* developed by Portik et al. 2017. It implements a scheme of sequential runs using the Nelder-Mead local optimization with initial random parameter values and perturbation of values between runs. We used the following settings of the dadi-pipeline tool: 3 rounds with 10, 20, 50 replications with 3-, 2-, and 1-fold perturbations, respectively, for each round. *dadi pipeline* was launched 50 times and the best resulting model had a log-likelihood equal to *−*1073.98. The average time for one launch of optimization was approximately 10 minutes.

To compare the runs with different initial demographic model structures (models 2 and 3), characteristics such as time for one iteration of the genetic algorithm, number of iterations, or mean and standard deviation of the log-likelihood value were calculated (Table 1). Launches with a simple initial structure show a more stable result in terms of the likelihood value but they have a longer average run time for one iteration. Furthermore, all the models obtained for simple-structure launches have the same demographic dynamics of population size change and similar parameters as final model reported in Gutenkunst et al. 2009, which is incorrect in cases involving launches of complex-structure models. At the same time, although the launches of models with complex structures result in a final model with a better log-likelihood score, it differs from the other models in terms of parameters values, which may indicate that it is inaccurate.

**FIG. 10.**
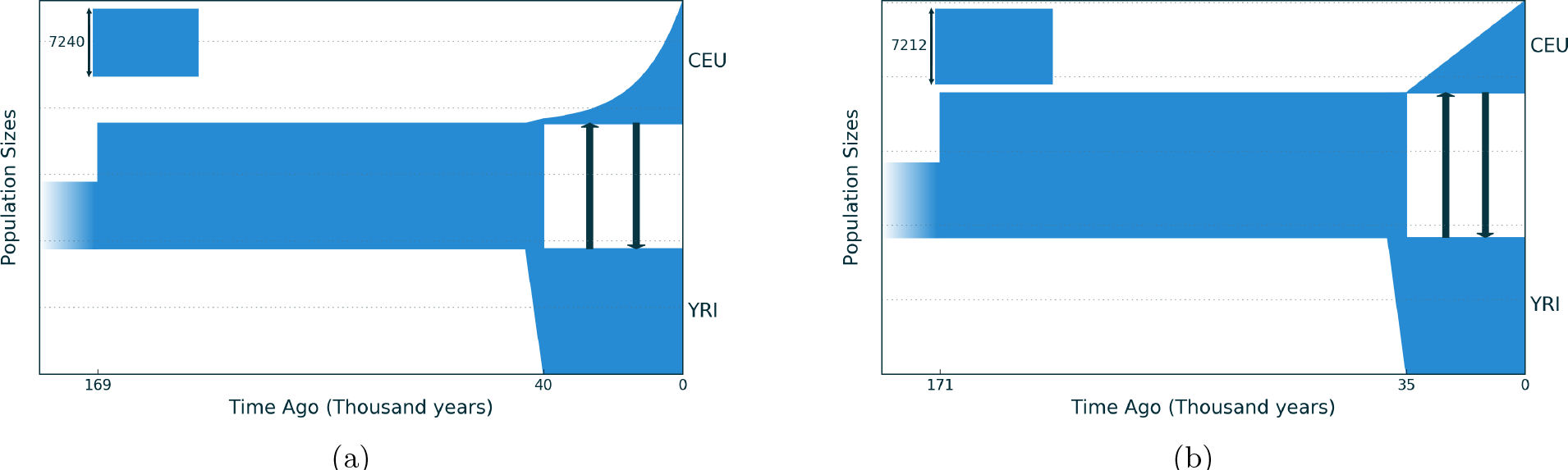
(a) Demographic model of two human populations (YRI and CEU) inferred using *∂*a*∂*i, as originally reported by Gutenkunst et al. 2009. This model has 6 parameters. The size of the African population after its growth 169,000 years ago is constant and migration rates between African and European populations are symmetric. (b) The demographic model from the same allele frequency spectrum inferred using GADMA with 12 parameters. Note that the differences between the inferred models is the slightly later age of the split between the YRI and CEU populations and the linear population growth (as opposed to exponential growth) in the CEU population in the model obtained with GADMA.

**Table 1.**
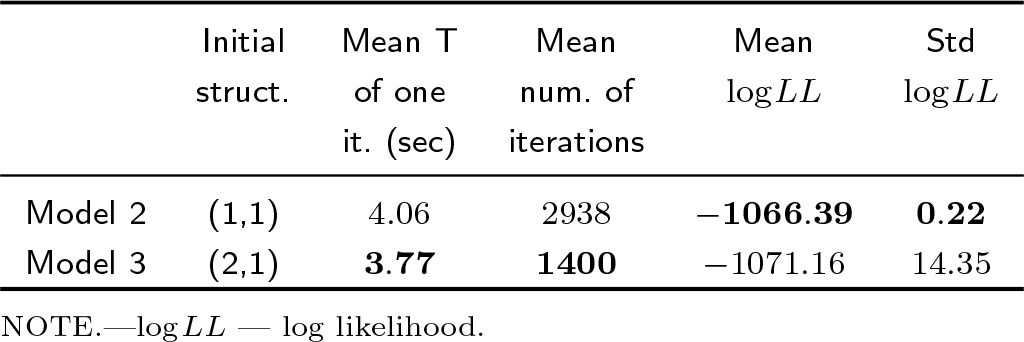
Comparison of runs using GADMA with different initial demographic model structures, simple (10 times) and complex (10 times), for the case of searching for the optimum demographic models of two human populations, YRI and CEU.

|  | Initial<br>struct. | Mean T<br>of one<br>it. (sec) | Mean<br>num. of<br>iterations | Mean<br>log LL | Std<br>log LL |
| --- | --- | --- | --- | --- | --- |
| Model 2 | (1,1) | 4.06 | 2938 | **-1066.39** | **0.22** |
| Model 3 | (2,1) | **3.77** | **1400** | -1071.16 | 14.35 |

NOTE.—log LL — log likelihood.

#### The YRI, CEU, CHB three population example

We also applied GADMA to infer demographic models for the case of three human populations based on same the AFS used in Gutenkunst et al. 2009. The first model (model 1) used the same parameters as in Gutenkunst et al. 2009, and their corresponding values were inferred (Table 4). GADMA yielded a better log-likelihood values for parameters than those reported in Gutenkunst et al. 2009. However, the timing of the split between the YRI population and the CEU+CHB populations was dated to 400,000 years ago, which is 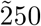 thousand years older than that inferred in any previous studies. To correct this, we restricted the age of this splitting event to 150,000 years ago based on previously published estimates (Cruciani *et al.*, 2011; Poznik *et al.*, 2013; Shriner *et al.*, 2014). A demographic model (model 2) was inferred with this age restriction, which yielded a better log-likelihood value than that in Gutenkunst et al. 2009. Next, we inferred the demographic model (model 3) that included all 26 parameters. Here we also observed an unrealistic earlier age for the divergence between YRI and CEU+CHB (results not shown). When we applied the 150,000 year age constraint as in model 2, we inferred a demographic model (model 3) that not only showed the highest log-likelihood value, but also the best *AIC* value.

**Table 2.** Maximum likelihood parameters for different demographic models for two human populations, YRI and CEU inferred using either *∂*a*∂*i or GADMA (the latter under three different parameter settings). The migration rates are per generation. 95% confidence intervals are indicated in brackets.

| | Model from<br>$\partial a \partial i$ example | Model from<br>GADMA (1) | Model from<br>GADMA (2) | Model from<br>GADMA (3) |
| --- | --- | --- | --- | --- |
| Number of parameters: | 6 | 6 | 12 | 12 |
| Log-likelihood: | -1066.35 | -1066.28 | -1065.87 | **-1065.15** |
| AIC score: | 2144.70 | **2144.56** | 2155.74 | 2154.30 |
| Parameters: |  |  |  |  |
| $N_A$ | 7240<br>[6940 – 7499] | 7230<br>[6925 – 7464] | 7200<br>[6917 – 7513] | 7210<br>[6967 – 7521] |
| $N_{AF0}$ | 13620<br>[13024 – 14115] | 13580<br>[13017 – 14134] | 13400<br>[10833 – 17965] | 14000<br>[11240 – 19332] |
| $N_{EU0}$ | 515<br>[439 – 595] | 530<br>[459 – 608] | 560<br>[401 – 663] | 25<br>[0 – 71] |
| $N_{EU}$ | 13360<sup>2</sup><br>[9935 – 16252] | 12400<sup>2</sup><br>[10445 – 14672] | 12100<sup>2</sup><br>[8462 – 18074] | 8950<sup>1</sup><br>[7196 – 10799] |
| $N_{AF}$ | (=$N_{AF0}$) | (=$N_{AF}$) | 13500<br>[11698 – 14834] | 13300<br>[11257 – 14350] |
| $m_{AF-EU} (\times 10^{-5})$ | 6.3<br>[5.4 – 7.4] | 6.3<br>[5.6 – 7.3] | 7.1<br>[10.3 – 18.7] | 5.8<br>[8.4 – 16.9] |
| $m_{EU-AF} (\times 10^{-5})$ | (=$m_{AF-EU}$) | (=$m_{AF-EU}$) | 6.1<br>[9.3 – 14.6] | 6.1<br>[9.3 – 14.4] |
| $T_{AF}$ (kya) | 168.5<br>[143.2 – 204.6] | 171.5<br>[143.7 – 204.8] | 176.5<br>[114.8 – 225.4] | 171.1<br>[118.1 – 203.6] |
| $T_{AF-EU}$ (kya) | 40<br>[36.7 – 43.6] | 40.8<br>[37.7 – 44.2] | 42.3<br>[34.9 – 46.5] | 34.8<br>[29.3 – 39.4] |

<sup>1</sup>Linear growth
<sup>2</sup>Exponential growth
NOTE.— $N_A$ — size of ancestral population; $N_{AF0}$ — size of ancestral population after growth; $N_{EU0}$ — size of European population after divergence of ancestral population; $N_{EU}$ — current size of European population (after growth); $N_{AF}$ — current size of African population; $m_{AF-EU}$ — migration rate from European pop. to African pop.; $m_{EU-AF}$ — migration rate from African pop. to European pop.; $T_{AF}$ — time of ancestral population size growth; $T_{AF-EU}$ — time of divergence.

**FIG. 11.**
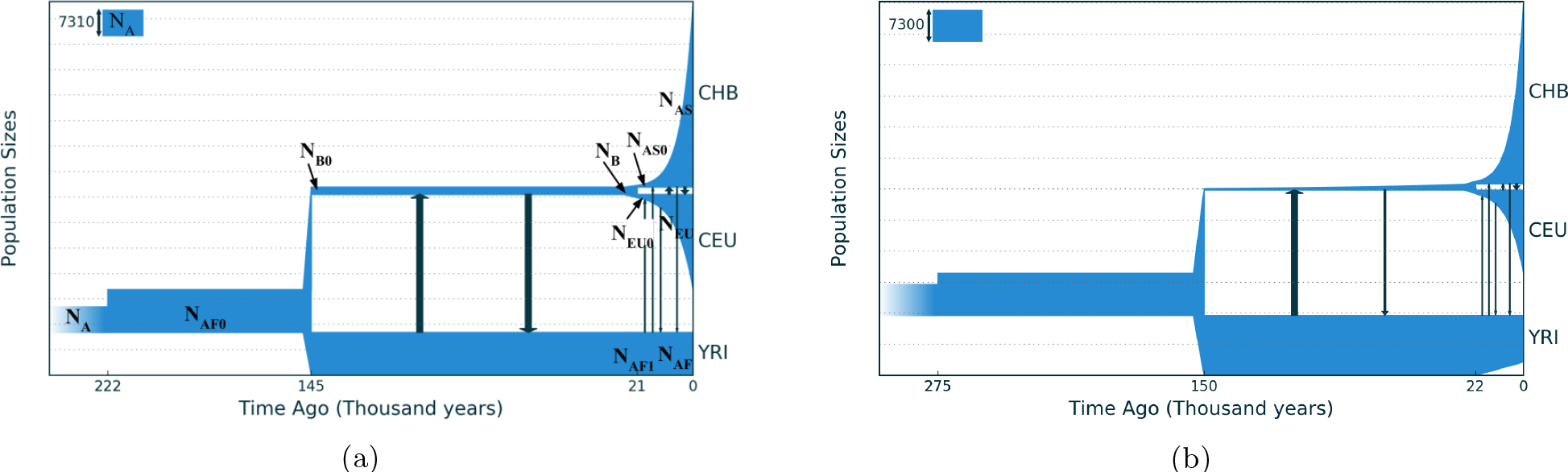
(a) Demographic model of three human populations (YRI, CEU, CHB) inferred using *∂*a*∂*i, as originally reported by Gutenkunst et al. 2009, which included 14 parameters. (b) The demographic model from the same allele frequency spectrum inferred using GADMA with 23 parameters.

**Table 3.** Comparison of outputs from different GADMA runs using either *∂*a*∂*i or *moments* for inferring demographic models for the YRI, CEU, CHB human populations with 26 parameters. For the runs performed with *∂*a*∂*i, the log-likelihood value was calculated using a *G* = {40,50,60} grid size.

| | Mean<br>time for one<br>iter. (sec) | Mean<br>num. of<br>iterations | Mean<br>$\log LL$ | Std<br>$\log LL$ |
| --- | --- | --- | --- | --- |
| $\partial a\partial i$ | 136.81 | 5531 | **-6293.56** | **7.26** |
| *moments* | **18.44** | **4700** | -6305.21 | 12.72 |

NOTE.— $\log LL$ — log-likelihood.

As in the case of testing two human populations, we also tested the three population case with the BFGS local optimization used in *∂*a*∂*i. We launched the optimization 100 times from randomly selected initial parameters and the best log-likelihood value obtained was −6323.99, which slightly differs from the optimal log-likelihood −6326.89.

With the exception of the earlier age of divergence between the YRI and CEU+CHB populations in model 1, demographic models 1 and 2 and the one inferred by Gutenkunst et al. 2009 have similar log-likelihood values and parameter estimates. In model 3, which has 26 parameters and the best *AIC* value, some parameters are also quite similar to the values in the other models. The major exceptions, however, are migration rates and the inferred population size of the Eurasian population, which exponentially grows from 200 to 1500 individuals after the split of the ancestral population. For comparison, in other models this number is a constant equal to 2,000 individuals, which seems less realistic than exponential growth. Migration rates vary considerably: they are higher compared to models 1 and 2 or the one found by Gutenkunst et al. 2009. The model shows that the largest migrations occurred between the YRI and CEU populations, and following the ancestral division, between the CEU and CHB populations. Moreover, the more geographically distant the populations are, the smaller is the observed migration rate.

GADMA was launched 10 times for each of the three models using *∂*a*∂*i and the best solutions were observed (Table 4). We also launched GADMA using *moments* (also 10 times) to compare its effectiveness with *∂*a*∂*i. The authors of *moments* conducted similar comparisons on simulated data (Jouganous *et al.*, 2017). Various characteristics of time and stability of the log-likelihood value based on the results of 20 GADMA runs (10 using *∂*a*∂*i, 10 using *moments*) are presented in the Table 3. Log-likelihood values of models inferred using *moments* were recalculated using *∂*a*∂*i with the *G* = {40,50,60} grid size simulated AFS so that log-likelihoods of the obtained with the two methods were comparable. Analyses using *moments* were 7.5 times faster than *∂*a*∂*i, whereas those using *∂*a*∂*i were more accurate: the average and variance of the likelihood values of the inferred models were less than the values inferred using *moments*.

**Table 4.** Maximum likelihood parameters inferred from the demographic models for the for YRI, CEU, CHB human populations. Migration rates are per generation. 95% confidence intervals are indicated in brackets.

|  | Model from<br>Gutenkunst et al. | Model from<br>GADMA (1) | Model from<br>GADMA (2) | Model from<br>GADMA (3) |
| --- | --- | --- | --- | --- |
| Param. number: | 13 | 13 | 14 | 26 |
| Max log-likelihood: | −6316.89 | −6314.40 | −6315.86 | −**6288.36** |
| AIC score: | 12659.78 | 12654.80 | 12657.72 | **12628.72** |
| Parameters: |  |  |  |  |
| $N_A$ | 7300<br>[6813–7480] | 6000<br>[5610–6030] | 7300<br>[6860–7476] | 7300<br>[6801–7588] |
| $N_{AF0}$ | 12300<br>[11197–12502] | 11840<br>[11268–12218] | 12200<br>[11148–12501] | 9900<br>[4789–20307] |
| $N_{B0}$ | 2100<br>[1982–2192] | 2050<br>[1960–2134] | 2070<br>[1981–2175] | 280<br>[13–224] |
| $N_B$ | ($=N_{B0}$) | ($=N_{B0}$) | ($=N_{B0}$) | 1450<sup>2</sup><br>[1315–1654] |
| $N_{AF1}$ | ($=N_{AF0}$) | ($=N_{AF0}$) | ($=N_{AF0}$) | 14000<br>[12011–14607] |
| $N_{AF}$ | ($=N_{AF0}$) | ($=N_{AF0}$) | ($=N_{AF0}$) | 11000<sup>1</sup><br>[8237–16299] |
| $N_{EU0}$ | 938<br>[769–986] | 930<br>[766–1037] | 950<br>[792–1018] | 890<br>[681–983] |
| $N_{EU}$ | 27300<sup>2</sup><br>[20537–32367] | 24700<sup>2</sup><br>[19707–31082] | 23700<sup>2</sup><br>[18630–30027] | 19600<sup>2</sup><br>[15658–26270] |
| $N_{AS0}$ | 510<br>[425–551] | 500<br>[444–565] | 510<br>[442–575] | 560<br>[493–679] |
| $N_{AS}$ | 53200<sup>2</sup><br>[39878–64334] | 50000<sup>2</sup><br>[38049–62437] | 46200<sup>2</sup><br>[37422–58526] | 42200<sup>2</sup><br>[26117–51110] |
| $m_{AF-B}$ ($\times 10^{-5}$) | 25<br>[23.7–27.5] | 26.2<br>[25.0–28.0] | 25.4<br>[24.1–27.6] | 17.0<br>[14.4–19.8] |
| $m_{B-AF}$ ($\times 10^{-5}$) | ($=m_{AF-B}$) | ($=m_{AF-B}$) | ($=m_{AF-B}$) | 57.4<br>[55.0–65.2] |
| $m_{AF-EU}$ ($\times 10^{-5}$) | 3.0<br>[2.34–3.9] | 3.1<br>[2.3–3.9] | 3.1<br>[2.5–4.1] | 4.5<br>[2.6–6.0] |
| $m_{EU-AF}$ ($\times 10^{-5}$) | ($=m_{AF-EU}$) | ($=m_{AF-EU}$) | ($=m_{AF-EU}$) | 2.8<br>[1.5–4.1] |
| $m_{AF-AS}$ ($\times 10^{-5}$) | 1.9<br>[1.1–2.7] | 1.9<br>[1.1–2.5] | 2.0<br>[1.2–2.6] | 0.9<br>[0.0–1.8] |
| $m_{AS-AF}$ ($\times 10^{-5}$) | ($=m_{AF-AS}$) | ($=m_{AF-AS}$) | ($=m_{AF-AS}$) | 2.1<br>[1.0–2.9] |
| $m_{EU-AS}$ ($\times 10^{-5}$) | 9.6<br>[7.9–12.2] | 10.2<br>[7.8–11.9] | 10.2<br>[8.2–12.3] | 18.8<br>[13.8–25.1] |
| $m_{AS-EU}$ ($\times 10^{-5}$) | ($=m_{AS-EU}$) | ($=m_{AS-EU}$) | ($=m_{AS-EU}$) | 6.8<br>[3.4–10.7] |
| $T_{AF}$ (kya) | 220.8<br>[170.0–590.1] | 570<br>[524.6–776.4] | 232<br>[165.9–617.3] | 274.8<br>[421.1–736.8] |
| $T_B$ (kya) | 144.0<br>[86.4–593.0] | 400<br>[331.3–638.8] | 150<br>[79.8–617.6] | 149.8<br>[135.3–324.4] |
| $T_{EU-AS}$ (kya) | 21.1<br>[19.9–21.9] | 21.0<br>[19.9–22.1] | 21.1<br>[20.1–22.1] | 22.4<br>[21.1–24.1] |

<sup>1</sup>Linear growth
<sup>2</sup>Exponential growth
NOTE.— $N_A$ — size of ancestral population; $N_{AF0}$ — size of ancestral population after growth; $N_{B0}$ — size of CEU+CHB population after divergence of ancestral population; $N_B$ — size of CEU+CHB population before its split; $N_{AF1}$ — size of YRI population at the time of CEU+CHB population divergence; $N_{AF}$ — current size of YRI population; $N_{EU0}$ — size of CEU population after divergence; $N_{EU}$ — current size of CEU population; $N_{AS0}$ — size of CHB population after divergence; $N_{AS}$ — current size of CHB population; $m_{1-2}$ — migration rate from population 2 to population 1; $T_{AF}$ — time of growth of size of ancestral population; $T_B$ — time of divergence of ancestral population to YRI population and CEU+CHB population; $T_{EU-AS}$ — time of divergence of CEU+CHB population to CEU population and CHB population.

### Demographic history of Gillittes checkerspot butterfly

We next tested GADMA using data from (McCoy *et al.*, 2014), which examined the demographic history of Gillettes checkerspot butterfly (*Euphydryas gillettii*). We used the same AFS as that used in the original paper, which consisted of eight individuals from the population in Colorado (CO) and eight individuals from the native population in Wyoming (WY). Four our analyses, we used two allele frequency spectra of size 13 × 13, one for synonymous SNVs only and one including all SNVs.

McCoy et al. tested three types of demographic models: 1) type A — models without migration between populations; 2) type B1 — models with unidirectional migration from CO to WY, and 3) type B2 — models with bidirectional migration between CO and WY. In the original paper, t the three demographic models were tested using the AFS based on synonymous SNVs only. Model A had the best *AIC* value, so the type A demographic model was inferred from the AFS using all SNVs. We used models A and B2 to infer the demographic models from both allele frequency spectra (synonymous SNVs and all SNVs) with GADMA.

Without considering migration, the models McCoy et al. 2013 had the following structure: there was one population of *N*_*A*_ size, which at some point of time was divided into two subpopulations, the size of which did not change in the future (sudden change of population size). All parameters were calculated with respect to *N*_*A*_ and had the following notation: *η*_*WY*_, *η*_*CO*_ relative population sizes at the current time; *τ*_*SPLIT*_ — time/age of population splitting event; and *M*_*WY*−*CO*_, *M*_*CO*−*WY*_ — scaled migration rates. For the models we inferred using GADMA, we also included the parameter *η*_*WY*0_ ∈ [0,1] — the size of the WY population immediately after the splitting event or the fraction of the size of the ancestral population that forms the WY population. The size of CO population immediately following the splitting event is equal to 1 − *η*_*WY*0_, because *N*_*A*_ = 1 before the division. However, in the case when the population size change is sudden, the size of the populations following the splitting event is equal to the size at the present time.

We ran four executions of GADMA with different data inputs: 1) the AFS using synonymous SNVs only without migration; 2) the AFS using synonymous SNVs only with migration; 3) the AFS using all SNVs without migration; and 4) the AFS using all SNVs with migration. For each execution, the analysis was repeated 50 times. *moments* was used to simulate the allele frequency spectrum due to its faster computational speed. However, the final log-likelihood scores that are presented in Tables S1 and 5 were calculated using *∂*a*∂*i with a *G* = {32,42,52} grid size, to compare our results with the original finding reported in McCoy et al. 2013.

**Table 5.**
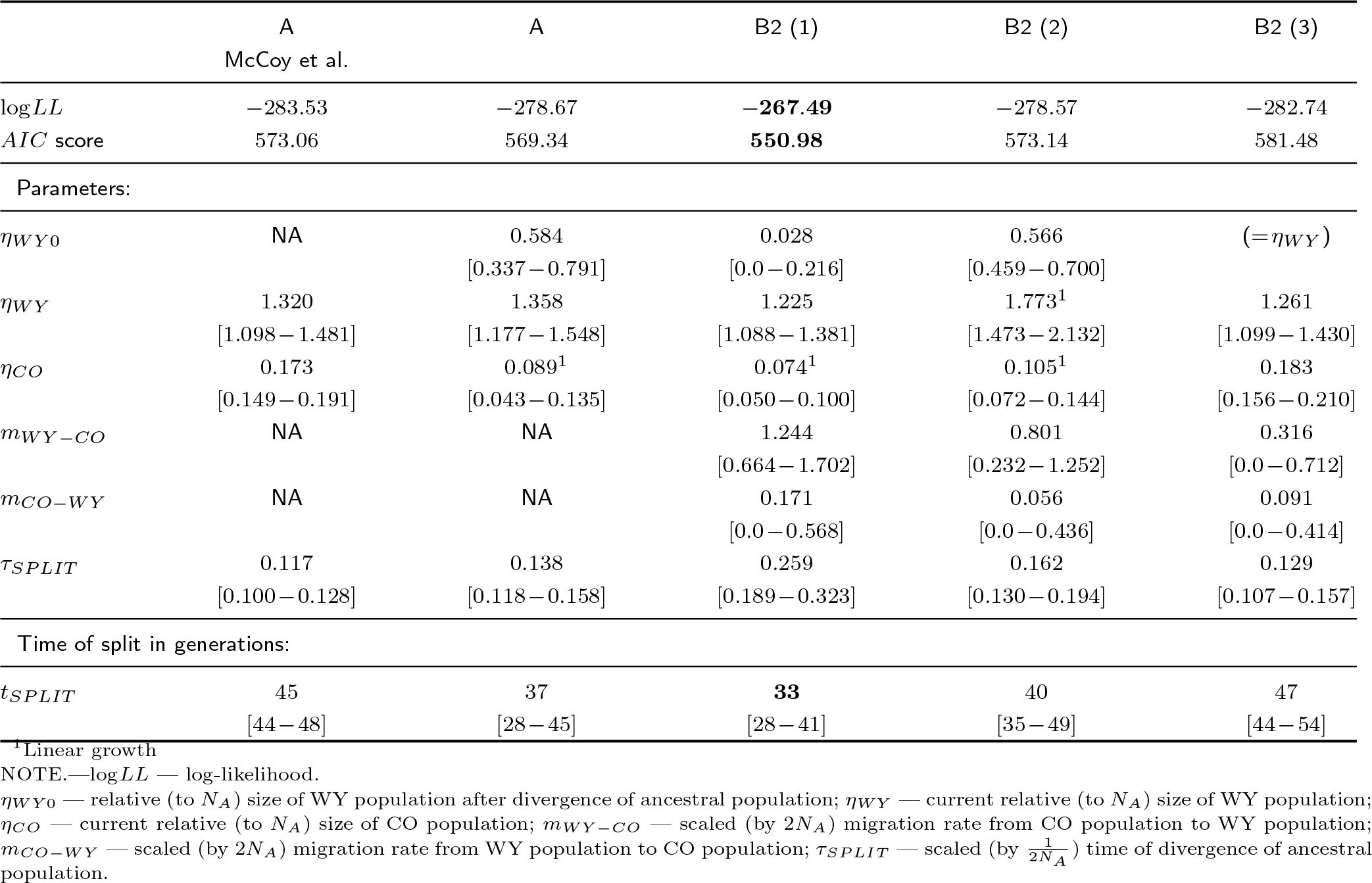
Demographic models and their associated parameters inferred in GADMA using an allele frequency spectrum based on all SNVs Gillettes checkerspot butterfly populations. 95% confidence intervals are indicated in brackets. The first column shows the parameters inferred in the original McCoy et al. 2013 paper.

The size of the generation of demographic models in the genetic algorithm was selected to be 10, the strength and mutation rate were set to 0.2, with constants 1.0 and 1.02, respectively, and the proportions of the best, mutated, crossed and random models in the new generation equal to 0.2:0.3:0.3:0.2. The likelihood was considered significant to two significant digits and the genetic algorithm stopped after 100 iterations without improvement. As for the local search, the Powell method was chosen. Since the structure of the demographic model from McCoy et al. 2013 corresponds to the simplest structure (1,1), it was chosen as both the initial and final structure of the demographic model.

Several runs of GADMA produced different local minima and the resulting alternative models and their inferred parameters are presented in Figure 12 and Table 5, respectively. We note that one of the models selected by GADMA, Model 26 (Figure 12d), was the same model as was inferred by McCoy et al. 2013, which includes a sudden growth of population size following the splitting of the ancestral population. For the AFS based synonymous SNVs, only models of type B2 had better likelihood values than models of type A, in contrast to the models inferred by McCoy et al. 2013.

**FIG. 12.**
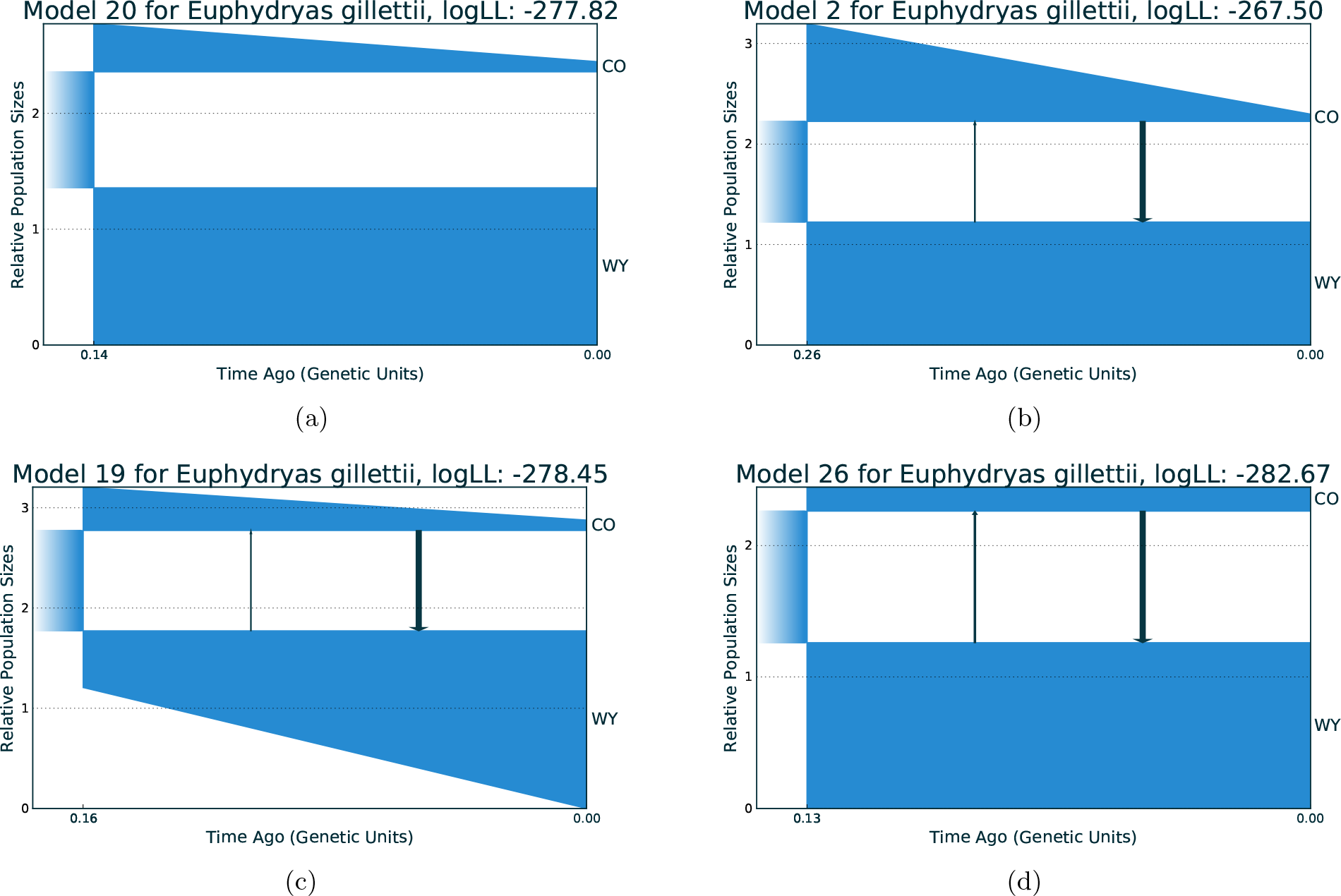
Alternative demographic models based on an AFS for Gillettes checkerspot butterflies from Colorado (CO) and Wyoming (WY), inferred with GADMA. Models of type B2 have negligible migration rates. (a) Type A model: without migration; (b) Type B2 model with migration; (c) Type B2 model with migration; (d) Type B2 model with migration.

One of the findings reported by McCoy et al. 2013 was that demographic models that include migration may not be applicable when the real population history shows no evidence for migration. This conclusion was based on two factors: 1) the type A demographic model, inferred from the AFS using synonymous SNVs, had the best *AIC* score; and 2) the estimate of the splitting time of the ancestral population in models that included migration had wide 95% confidence intervals, such that the parameter boundaries included zero (see McCoy et al. 2013). However, during our regeneration of these authors results, we found errors associated with some of the parameters of the demographic models: split time of the ancestral population and migrations were mixed up with each other, resulting in migrations, but not split times, having wide confidence intervals that include zero. Moreover, our analyses using GADMA showed that the 95

The average population size of butterflies from Colorado (CO) was estimated as *N*_*CO*_ = 34 individuals by McCoy et al. 2013. If we scale the parameters of the best-fitting model so that the average size of CO is equal to this value (*x*·(*η*_*CO*_ +(1 − *η*_*WY*0_))/2 = *N*_*CO*_), then we get 33.6 generations (*t*_*SPLIT*_ = 2·*x*·*τ*_*SPLIT*_) after the division of the ancestral population, which corresponds to the actual 33 generations observed (1977-2010). Such estimates were made for all resulting models (Table 5). However, the best-fitting model had the best estimated value of split time of the ancestral population (33 generations).

### Demographic history of the Gaboon forest frog

For our third evaluation of performance, we compared the demographic models inferred using GADMA with those inferred with the recently developed optimization method implemented in the *dadi pipeline* (Portik *et al.*, 2017). This pipeline uses *∂*a*∂*i to simulate an AFS and infers parameters of the researcher-specified demographic model by several rounds of consistent runs using the Nelder-Mead local optimization method. During the first round, random values are estimated and during each successive round, the best parameters from the previous round are used. Prior to each Nelder-Mead local optimization, current parameter values are perturbed. Portik et al., 2017 demonstrated the *dadi pipeline* using allele frequency spectra generated from the Gaboon forest frog (*Scotobleps gabonicus*) and these same data was used to perform the analyses using GADMA. Sampling included eighty-four samples from 33 localities of Lower Guinea, West Africa, which were divided into Northern and Southern populations according to hierarchical Bayesian clustering analysis of 7,633 unlinked SNPs. Each Northern and Southern group was further divided into three clusters: Cameroon Volcanic Line North (CVLN) and Cameroon Volcanic Line South (CVLS) and Cross River (CrossRiver) populations for the Northern group, and North Coast, South Coast and East Gabon populations for the Southern group.

In order to perform our analyses using GADMA, three two-dimensional folded allele frequency spectra were chosen from Portik et al., 2017: 1) a 41 × 19 spectrum for Northern and Southern populations; 2) a 31 × 19 spectrum for CVLN and CVLS; and 3) a 15 × 31 for the CrossRiver and CVLN populations. In generating the allele frequency spectra, only a single SNP per RAD locus was kept, so loci are assumed to be independent. For each spectrum, we estimated best-fitting log-likelihoods, *AIC* scores, Akaike weights (*ω*_*i*_) (Burnham and Anderson, 2003) and parameters for 14 demographic models. Two demographic models assumed no divergence between populations, while the remaining 12 models assumed a split of the ancestral population and different assumptions related to migration rates, isolation, and population size changes. All population size changes were considered to have sudden change dynamics, which we included in our analyses as well. For all models, we provide inferred parameter values in Tables S2, S3, and S4, including value of *θ* = 4*N*_*A*_*μL*, where *μ* is the mutation rate per generation per site and *L* is effective sequence length.

The best-fitting demographic model inferred using the *dadi pipeline* for the Northern and Southern populations only suggests a population expansion, followed by secondary contact and symmetrical migration between populations (Δ*AIC* = 13.6, *ω*_*i*_ = 0.99). For the CVLN and CVLS populations, the best demographic model included population expansion, secondary contact and asymmetrical gene flow from CVLN to CVLS (Δ*AIC* = 94.2, *ω*_*i*_ = 0.99). Finally, two demographic models explain the AFS data equally well for the CrossRiver and CVLN populations: one with secondary contact but no population size change and asymmetrical migration from CVLN to Cross River (*ω*_*i*_ = 0.53) and another that included an ancestral population division with continuous asymmetric gene flow (*ω*_*i*_ = 0.43).

We inferred all 12 demographic models with divergence as in Portik et al. 2017 for each of the three allele frequency spectra with GADMA. Each of 36 runs was repeated 10 times as opposed to the 50 runs launched with the the *dadi pipeline*. We originally had planned to conduct the analyses with *∂*a*∂*i for GADMA optimization to simulate the allele frequency spectra with the same grid size as that used by Portik et al. 2017: *G* = {40,50,60} for the CrossRiver, CVLN and CVLN, CVLS populations; and *G* = {50,60,70} for the Northern and Southern populations. However, we found that the optimization with *∂*a*∂*i was unstable and therefore, we set upper bounds on population size (12*N*_*ref*_) and generation time (5*N*_*ref*_). Under these parameters, *∂*a*∂*i was found to be stable, but some parameters were equal to the upper bounds and in such cases GADMA was launched 10 times for each model using *moments* with larger upper bounds (100*N*_*ref*_ for population size and 10*N*_*ref*_ for generation time). The inferred demographic models and their estimated parameters are presented in Tables S2-S4. We report demographic models estimated with *moments*, and if a similar model was inferred by both *∂*a*∂*i and *moments*, we only report the result with the best log-likelihood value.

**Table 6.** Demographic models and associated parameters inferred in GADMA for the Northern and Southern populations of the Gaboon forest frog. LogLL — log-likelihood. Expected AFS was simulated using *∂*a*∂*i with *G* = {50,60,70}. See text for more details.

|  | Best<br>from<br>dadi pipeline<sup>1</sup> | Model,<br>structure<br>(1,2) | Best<br>from<br>GADMA<sup>2</sup> |
| --- | --- | --- | --- |
| Number of parameters | 7 | 11 | 9 |
| Log-likelihood | -439.91 | -402.20 | **-402.00** |
| AIC score | 893.81 | 826.40 | **821.99** |
| Parameters: |  |  |  |
| $\theta$ | 573.9 | 134.7 | 107.0 |
| $\nu_1^a$ | 0.259 | 1.894 | 2.515 |
| $\nu_2^a$ | 0.247 | 2.634 | 3.339 |
| $m_{12}^a$ | NA | 0.046 | 0.033 |
| $m_{21}^a$ | NA | 0.0 | NA |
| $\nu_1^b$ | 3.121 | 13.544 | 17.139 |
| $\nu_2^b$ | 1.291 | 5.584 | 7.057 |
| $m_{12}^b$ | 0.049 | 0.008 | 0.006 |
| $m_{21}^b$ | $=m_{12}^b$ | 0.017 | 0.014 |
| $T_a$ | 0.302 | 4.335 | 5.779 |
| $T_b$ | 0.556 | 2.162 | 2.692 |

<sup>1</sup> Divergence, secondary contact with population size change and symmetric migration.
<sup>2</sup> Divergence, unidirectional migration from Southern to Northern, followed by population size growth and bidirectional asymmetric migrations.
NOTE.— $\theta=4N_A\mu L$ ; $\nu_p^x$ — relative (to $N_A$ ) size of population $p$ (1 for Northern, 2 for Southern) during time interval $x$ after divergence ( $a$ — first time interval, $b$ — second time interval); $m_{pk}^x$ — scaled (by $2N_A$ ) migration rate from population $k$ to population $p$ during time interval $x$ ; $T_x$ — scaled (by $\frac{1}{2N_A}$ ) time of time interval $x$ .

**FIG. 13.**
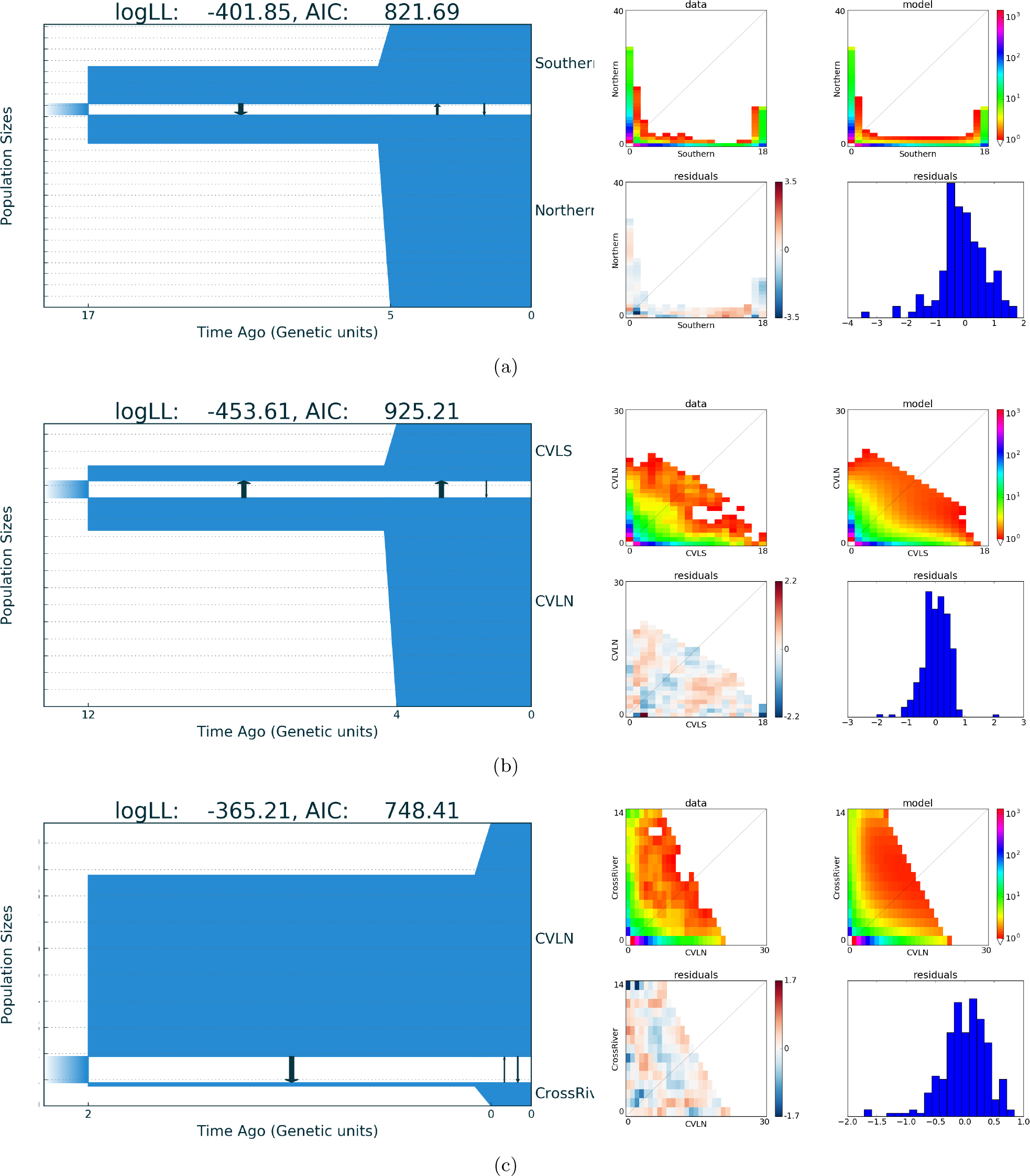
Demographic models with the highest log-likelihoods obtained from allele frequency spectra for different populations of the Gaboon forest frog, inferred using GADMA. (a) Demographic model inferred for the Northern, Southern populations;(b) Demographic model inferred for the CVLN and CVLS populations; and (c) Demographic model inferred for the CrossRiver and CVLN populations. The plots to the right of the demographic models show the AFS data (upper-left), AFS of the demographic model (upper-right), Anscombe residuals between model and data (lower-left) and the histigram of the Anscombe residuals (lower-right).

For almost all (32 from 36) cases, GADMA was able to infer better models in terms of both log-likelihood scores and parameters than were previously inferred using the *dadi pipeline* in Portik et al. 2017. Moreover, we obtained better alternative models for the CrossRiver, CVLN, Northern, and Southern populations. These models included an initial split of the ancestral population with ancestral asymmetric migration and a population size change (Δ*AIC* = 3.24, *ω*_*i*_ = 0.82) for the CrossRiver and CVLN populations; and secondary contact with asymmetric migration and population size change (Δ*AIC* = 6.82, *ω*_*i*_ = 0.97) for the Northern and Southern populations. For the third AFS of the CVLN and CVLS populations, the model that includes secondary contact with asymmetric migration and population size change was found to be superior based on comparison of the Akaike information criterion (Δ*AIC* = 87.98, *ω*_*i*_ > 0.99), similar to the findings by Portik et al., but it received higher log-likelihood value: −455.17 using GADMA versus −463.3 in Portik et al., 2017.

**Table 7.** Demographic models and associated parameters inferred in GADMA for the CVLN and CVLS populations of the Gaboon forest frog. LogLL — log-likelihood. Expected AFS was simulated using *∂*a*∂*i with *G* = {40,50,60}. See text for more details.

|  | Best<br>from<br>dadi pipeline | Model,<br>structure<br>(1,2) | Best<br>from<br>GADMA |
| --- | --- | --- | --- |
| Number of parameters | 8 | 11 | 9 |
| Log-likelihood | -463.33 | -453.67 | **-453.65** |
| *AIC* score | 942.67 | **929.34** | **925.30** |
| Parameters: |  |  |  |
| $\theta$ | 572.18 | 134.17 | 145.5 |
| $\nu_1^a$ | 0.338 | 2.010 | 1.893 |
| $\nu_2^a$ | 1.331 | 0.840 | 0.880 |
| $m_{12}^a$ | NA | 0.0 | NA |
| $m_{21}^a$ | NA | 0.400 | 0.399 |
| $\nu_1^b$ | 4.528 | 13.220 | 12.266 |
| $\nu_2^b$ | 1.193 | 3.620 | 3.349 |
| $m_{12}^b$ | 0.108 | 0.0419 | 0.046 |
| $m_{21}^b$ | 1.084 | 0.365 | 0.395 |
| $T_a$ | 0.325 | 5.000 | 4.148 |
| $T_b$ | 0.870 | 1.970 | 1.808 |

<sup>1</sup> Divergence, secondary contact with population size change and asymmetric migration.
<sup>2</sup> Divergence, unidirectional migration from CVLN to CVLS, followed by population size growth and bidirectional asymmetric migrations.
NOTE.— $\theta = 4N_A\mu L$ ; $\nu_p^x$ — relative (to $N_A$ ) size of population $p$ (1 for CVLN, 2 for CVLS) during time interval $x$ after divergence ( $a$ — first time interval, $b$ — second time interval); $m_{pk}^x$ — scaled (by $2N_A$ ) migration rate from population $k$ to population $p$ during time interval $x$ ; $T_x$ — scaled (by $\frac{1}{2N_A}$ ) time of time interval $x$ .

**Table 8.** Demographic models and associated parameters inferred in GADMA for the CrossRiver and CVLN populations of the Gaboon forest frog. LogLL — log-likelihood. Expected AFS was simulated using *∂*a*∂*i with *G* = {40,50,60}. See text for more details.

|  | Best<br>from<br>dadi pipeline | Model,<br>structure<br>(1,2) | Best<br>from<br>GADMA |
| --- | --- | --- | --- |
| Number of parameters | 6 | 11 |  |
| Log-likelihoods | -377.99 | **-365.26** | -365.29 |
| *AIC* score | 769.98 | 752.52 | **748.58** |
| Parameters: |  |  |  |
| $\theta$ | 287.96 | 250.58 | 251.88 |
| $\nu_1^a$ | 0.350 | 0.149 | 0.139 |
| $\nu_2^a$ | 6.746 | 7.034 | 6.899 |
| $m_{12}^a$ | NA | 2.510 | 2.639 |
| $m_{21}^a$ | NA | 0.001 | NA |
| $\nu_1^b$ | $= \nu_1^a$ | 0.974 | 0.889 |
| $\nu_2^b$ | $= \nu_2^a$ | 8.707 | 8.873 |
| $m_{12}^b$ | 1.272 | 0.507 | 0.556 |
| $m_{21}^b$ | 0.221 | 0.328 | 0.312 |
| $T_a$ | 0.495 | 1.110 | 1.089 |
| $T_b$ | 0.402 | 0.101 | 0.109 |

<sup>1</sup> Divergence, secondary contact with no population change and asymmetric migration.
<sup>2</sup> Divergence, unidirectional migration from CVLN to CrossRiver, followed by population size growth and bidirectional asymmetric migrations.
NOTE.— $\theta = 4N_A\mu L$ ; $\nu_p^x$ — relative (to $N_A$ ) size of population $p$ (1 for CrossRiver, 2 for CVLN) during time interval $x$ after divergence ( $a$ — first time interval, $b$ — second time interval); $m_{pk}^x$ — scaled (by $2N_A$ ) migration rate from population $k$ to population $p$ during time interval $x$ ; $T_x$ — scaled (by $\frac{1}{2N_A}$ ) time of time interval $x$ .

We next inferred the demographic model with a structure equal to (1,2) with GADMA (Tables 6, 7 and 8). For all three allele frequency spectra, such a model had the best log-likelihood among previously inferred models (Tables S2, S3 and S4). For the Northern and Southern populations, the demographic model with the (1,2) structure also had the best *AIC* score. This model showed similar features for all observed data: an interval of time after the splitting of the ancestral population with unidirectional migration, followed by a time interval with bidirectional migration and population size change. The direction of the unidirectional migration was as follows: from CVLN to CrossRiver; from CVLN to CVLS; and from Southern to Northern. We also constructed two additional demographic models based on observed features and inferred their parameters: splitting of the ancestral population with unidirectional migration followed by either symmetric or asymmetric migrations with population size change. The demographic model with unidirectional migration followed by asymmetric migration and population size change was found to be the best for all three allele frequency spectra (Tables 6, 7 and 8): Northern and Southern populations: Δ*AIC* = 4.41, *ω*_*i*_ = 0.89; CVLN and CVLS populations: Δ*AIC* = 1.04, *ω*_*i*_ = 0.58; and CrossRiver and CVLN populations: Δ*AIC* = 0.04, *ω*_*i*_ = 0.44. But there were several additional demographic models that explained the data equally well: the same demographic model as just described but which also included symmetric migration for CrossRiver, CVLN (*ω*_*i*_ = 0.43) and secondary contact with asymmetric migration and population size change for CVLN and CVLS (*ω*_*i*_ = 0.34).

## Discussion

We report the development and mathematical justification of GADMA and demonstrate its effectiveness using several previously published data sets. GADMA is the first program that allows the automatic inference of demographic history of up to three populations from an allele frequency spectrum. Our method is based on the genetic algorithm and uses existing solutions from either *∂*a*∂*i or *moments* to simulate the AFS from the proposed demographic model. Existing optimizations, implemented in either *∂*a*∂*i and *moments*, require prior specification of demographic models to be tested and are thus inefficient in practice, given the large number of possible demographic scenarios that can be constructed for one or more populations. Our method is implemented in the GADMA software, which is openly accessible via the link https://github.com/ctlab/GADMA.

GADMA was shown to be efficient in performance: it was applied to three different datasets, representing three different organismal systems and associated demographic histories and the inferred demographic models had better log-likelihood scores than those reported in the original papers, which were derived from optimizations using either *∂*a*∂*i or moments alone. Moreover, the demographic histories inferred with GADMA were consistent with the known history of the three taxa (Gutenkunst *et al.*, 2009; McCoy *et al.*, 2014; Poznik *et al.*, 2013). We also demonstrated the stability of the search, starting with demographic models with simpler structures rather than more complex ones, which reflects the profitability of using a search scheme that includes an increase in model structure complexity. Additionally, we compared pipelines using *moments* or *∂*a*∂*i and showed that the computational speed of *moments* was much greater than for *∂*a*∂*i. Thus, GADMA is the first software that effectively infers a demographic model from an allele frequency spectrum with nothing required from the user, except the structure of the demographic model, which determines the extent of the model complexity and associated details.

Unfortunately, there are some limitations on how informative the AFS can be with regards to historical demographic inference. For example, it has been shown that the AFS of a single panmictic population can be matched to different demographic scenarios (Myers *et al.*, 2008). We should expect the same behavior in the case of multiple populations, which requires estimating the joint AFS. This problem can be solved by using some additional information about observed populations, for example, two-locus statistics (Ragsdale and Gutenkunst, 2017). Incorporating genetic linkage information with AFS data could also improve the accuracy of inference of demographic history.

In our analyses using the AFS from three human populations (YRI, CEU and CHB), we inferred a best-fitting demographic model that showed an expansion out of Africa around 400 thousand years ago, which is not supported by previous studies (Gutenkunst *et al.*, 2009; Scheinfeldt *et al.*, 2010; Shriner *et al.*, 2016). This can be caused by the limitation of the informativeness of the AFS or by noise in the spectrum as a result of including low-quality variant calls. However, Gutenkunst et al. 2009 noted the high quality of this data set. The demographic model inferred with GADMA using the same parameters as in Gutenkunst et al. 2009, under the assumption that expansion took place not earlier than 150 thousand years ago, resulted in parameter values similar to those reported in Gutenkunst et al. 2009. We also inferred all possible parameters, including asymmetric migration rates and different dynamics of population size changes, and obtained a demographic model with the best *AIC* score. With GADMA, we observed higher asymmetric migration rates and the growth of the Eurasian population after its split from the African population.

Allele frequency spectra of two isolated populations of Gillittes checkerspot buttery from Wyoming and Colorado showed several alternative models with values very close to the best value of the composite likelihood. All migrations that were inferred are negligible, which confirmed the isolation of the two populations. One of the inferred models is consistent with the demographic history that was estimated previously by (McCoy *et al.*, 2014). The demographic model inferred using GADMA with the best likelihood value seems to be a better model overall because, in addition to the best *AIC* score, it correctly inferred the timing of the population split to the actual known value of 3̃3 years. However, we suggest that further research is necessary to identify alternative models that may better fit the demographic history of these checkerspot butterfly populations.

We conducted a series of experiments for selecting demographic models for the Gaboon forest frog, also repeating the analyses performed by (Portik *et al.*, 2017). Nearly all of the 12 models inferred previously were found to be suboptimal. For two of the three populations sets analyzed with GADMA, demographic models with higher log-likelihoods were chosen compared to those previously inferred. For the comparison that included the CVLN and CVLS populations, the demographic model with the highest log-likelihood was consistent with model inferred by Portik et al., 2017, but new values of parameters with better likelihood values were found. Demographic model optimization using *∂*a*∂*i proved to be unstable, and we were consequently forced to either specify lower limits on specific parameter values or instead use *moments* for model optimization. *moments* proved to be indeed more stable in simulating the expected AFS from the demographic model. We then inferred the full parameters of the demographic model with a structure equal to (1,2) for each of three datasets. For Northern and Southern populations, models using this structure resulted in higher log-likelihood scores. However, we noticed some peculiar properties in parameter values and generated new demographic models based on these properties and inferred their parameters using the three allele frequency spectra. These analyses resulted in a model with improved *AIC* scores for each of the three allele frequency spectra. This model contains divergence of the ancestral population, a time interval with unidirectional migration followed by a time interval with population size change and bidirectional asymmetric gene flow.

While the optimization search implemented in GADMA is able to find demographic models with the best likelihood score and their associated parameters, it is important to minimize the number of parameters so as to avoid the possibility of overfitting the model to the AFS data. Fortunately, such a strategy is included in in GADMA using *AIC* scores: it informs the user about overfitting when the demographic model with best likelihood score and best *AIC* score do not match. Additionally, GADMA can infer demographic models with all possible parameters, allowing researchers to explore additional models based on the inferred model, as we demonstrated in the case of the Gaboon forest frog. However, we note that GADMA it does not sort out all possible number of parameters, so it is not guaranteed to find model with best *AIC* score.

Another direction in the further development of our work is increasing the number of considered populations. Currently, GADMA can analyze up to three populations, similar to *∂*a*∂*i. In contrast, *moments* can simulate allele frequency spectra for up to five populations. Moreover, including the estimation of selection coefficients (e.g., Gutenkunst et al. 2009) and the development of a user-friendly interface for various types of data sets (e.g., all SNPs, synonymous SNPs only, etc.), will help to further expand the capabilities of GADMA. It is also possible to improve the proposed method, using various modifications of the genetic algorithm, for example, one that infers deliberately different demographic models (Squillero and Tonda, 2016).

## Availability of source code and requirements

- Project name: GADMA
- Project home page: https://github.com/ctlab/GADMA
- Operating system(s): Platform independent
- Programming language: Python
- Other requirements:

- Python (2.5, 2.6, 2.7),
- NumPy (≥ 1.2.0),
- Scipy (≥ 0.6.0),
- *∂*a*∂*i (≥ 1.7.0) or *moments* (≥ 1.0.0)
- License: GNU GPL v3.

## Availability of supporting data and materials

All data, parameters for GADMA runs and results are available in the supplementary materials.

## Supporting information

All data, parameters for GADMA runs and results

## Acknowledgements

We would like to express our sincere thanks to Maxim Buzdalov for the fruitful discussion about genetic algorithm, Victoria Efremova for the proofreading of GADMA manual, Nikita Alexeev for the statistical assistance, Rajiv Mccoy and Daniel Portik for the provided AFS data and useful discussions about their papers. This work was supported by the JetBrains Research.

