## Supplemental Tables S1-S4 for "GADMA: Genetic algorithm for inferring demographic history of multiple populations from allele frequency spectrum data"

November 2018

Table S1: Demographic models for allele frequency spectrum using synonymous SNVs only of Butterfly *Euphydryas gillettii* populations (WY — Wyoming, CO — Colorado). 95% confidence intervals are indicated in brackets.

| | A [1] | B1 [1] | B2 [1] | A | B2 (1) | B2 (2) | B2 (3) |
|---|---|---|---|---|---|---|---|
| Number of parameters: | 3 | 4 | 5 | 4 | 6 | 6 | 6 |
| $\log LL$ | −211.63 | −210.83 | −210.80 | −211.42 | **−205.94** | −210.59 | −210.80 |
| AIC | 429.26 | 429.66 | 431.60 | 430.84 | **423.88** | 433.18 | 433.60 |
| Parameters: | | | | | | | |
| $\eta_{WY0}$ | NA | NA | NA | 0.79 | 0.002 | 0.999 | $(=\eta_{WY})$ |
| | | | | [0.540 − 0.953] | [0.0 − 0.275] | [0.888 − 1.076] | |
| $\eta_{WY}$ | 0.922 | 0.884 | 0.893 | 0.960 | 0.856 | 0.742[1] | 0.873 |
| | [0.670 − 1.203] | [0.630 − 1.126] | [0.655 − 1.076] | [0.690 − 1.056] | [0.690 − 1.056] | [0.384 − 1.113] | [0.637 − 1.123] |
| $\eta_{CO}$ | 0.104 | 0.119 | 0.121 | 0.064[1] | 0.048[1] | 0.120 | 0.121 |
| | [0.079 − 0.135] | [0.093 − 0.155] | [0.081 − 0.166] | [0.012 − 0.089] | [0.012 − 0.089] | [0.090 − 0.162] | [0.086 − 0.166] |
| $m_{WY-CO}$ | NA | 0.887 | 0.906 | NA | 2.857 | 0.952 | 0.923 |
| | | [0.0 − 1.889] | [0.016 − 1.933] | | [1.653 − 3.763] | [0.316 − 1.916] | [0.125 − 2.050] |
| $m_{CO-WY}$ | NA | NA | 0.002 | NA | 0.176 | 0.0 | 0.0 |
| | | | [0.0 − 0.634] | | [0.0 − 1.281] | [0.0 − 0.364] | [0.0 − 0.379] |
| $\tau_{SPLIT}$ | 0.066 | 0.080 | 0.081 | 0.075 | 0.241 | 0.079 | 0.080 |
| | [0.049 − 0.086] | [0.059 − 0.107] | [0.050 − 0.118] | [0.132 − 0.368] | [0.132 − 0.368] | [0.058 − 0.112] | [0.054 − 0.118] |

[1] Linear growth

Descriptions of model names in tables for Gaboon forest frog *Scotobleps gabonicus*

| | |
|---|---|
| anc_asym_mig | Split, Ancient Asymmetrical Migration. |
| anc_asym_mig_size | Split, Ancient Asymmetrical Migration, Size Change. |
| anc_sym_mig | Split, Ancient Symmetrical Migration. |
| anc_sym_mig_size | Split, Ancient Symmetrical Migration, Size Change. |
| asym_mig | Asymmetrical Gene Flow. |
| no_mig | Split with No Migration. |
| no_mig_size | Split with No Migration, Size Change. |
| sec_contact_asym_mig | Split, Secondary Contact, Asymmetrical Migration. |
| sec_contact_asym_mig_size | Split, Secondary Contact, Asymmetrical Migration, Size Change. |
| sec_contact_sym_mig | |
| sec_contact_sym_mig_size | Split, Secondary Contact, Symmetrical Migration |
| Structure 1,2 | Split, Two Time Intervals with Asymmetrical Migrations. |
| sym_mig | Split, Symmetrical Gene Flow. |
| unidir_asym_mig_size | Split, Unidirectional Gene Flow followed by Asymmetrical Bidirectional Migrations and Size Change. |
| unidir_sym_mig_size | Split, Unidirectional Gene Flow followed by Symmetrical Bidirectional Migrations and Size Change. |

Table S2: Demographic models for Northern, Southern populations of Gaboon forest frog *Scotobleps gabonicus*. Expected likelihood was received by using of ∂a∂i with $G = [50, 60, 70]$ grid size.

| | N | Prev. $\log LL$ | Prev. $AIC$ | $\log LL$ | $AIC$ | $\Delta AIC$ | $\omega_i$ | $\theta$ | $\nu_1^a$ | $\nu_2^a$ | $\nu_1^b$ | $\nu_2^b$ | $m_{12}^a$ | $m_{21}^a$ | $m_{12}^b$ | $m_{21}^b$ | $T_a$ | $T_b$ |
|---|---|---|---|---|---|---|---|---|---|---|---|---|---|---|---|---|---|---|
| unidir_asym_mig_size | 9 | - | - | -402.00 | 821.99 | 0.00 | 0.89 | 107.0 | 2.515 | 3.339 | 17.139 | 7.057 | 0.033 | - | 0.006 | 0.014 | 5.779 | 2.692 |
| Structure 1,2 | 11 | - | - | -402.20 | 826.40 | 4.41 | 0.10 | 134.7 | 1.894 | 2.634 | 13.544 | 5.584 | 0.046 | 0.000 | 0.008 | 0.017 | 4.335 | 2.162 |
| sec_contact_asym_mig_size | 8 | -445.7 | 907.4 | **-407.21** | 830.42 | 8.43 | 0.01 | 262.6 | 1.128 | 1.246 | 6.972 | 2.906 | - | - | 0.019 | 0.033 | 1.621 | 1.091 |
| unidir_sym_mig_size | 8 | - | - | -410.26 | 836.52 | 14.53 | 0.00 | 254.3 | 0.987 | 1.304 | 7.045 | 3.043 | 0.055 | - | 0.021 | $m_{12}^b$ | 1.721 | 1.181 |
| sec_contact_sym_mig_size | 7 | -439.9 | 893.8 | **-411.62** | 837.24 | 15.25 | 0.00 | 279.5 | 1.024 | 1.190 | 6.471 | 2.783 | - | - | 0.025 | $m_{12}^b$ | 1.445 | 1.032 |
| anc_asym_mig_size* | 8 | -522.5 | 1061.0 | **-499.19** | 1014.38 | 192.39 | 0.00 | 90.2 | 3.885 | 5.161 | 24.025 | 8.599 | 0.048 | 0.020 | - | - | 8.579 | 2.170 |
| anc_sym_mig_size | 7 | -509.8 | 1033.6 | **-501.56** | 1017.12 | - | - | 80.2 | 5.182 | 5.228 | 27.808 | 10.110 | 0.028 | $m_{12}^a$ | - | - | 10.000 | 2.218 |
| anc_asym_mig_size | 8 | -522.5 | 1061.0 | **-500.62** | 1017.24 | 195.25 | 0.00 | 178.2 | 1.873 | 2.543 | 12 | 4.338 | 0.100 | 0.039 | - | - | 3.780 | 1.145 |
| anc_sym_mig_size | 7 | -509.8 | 1033.6 | **-503.66** | 1021.32 | 199.33 | 0.00 | 181.2 | 2.145 | 2.220 | 12 | 4.450 | 0.068 | $m_{12}^a$ | - | - | 3.806 | 1.051 |
| no_mig_size | 5 | -570.3 | 1150.6 | **-569.89** | 1149.78 | 327.79 | 0.00 | 570.1 | 0.115 | 0.139 | 3.178 | 1.340 | - | - | - | - | 0.085 | 0.588 |
| sec_contact_asym_mig | 6 | -674.6 | 1361.2 | **-643.79** | 1299.58 | 477.59 | 0.00 | 209.4 | 6.061 | 2.631 | $\nu_1^a$ | $\nu_2^a$ | - | - | 0.036 | 0.063 | 3.441 | 0.218 |
| sec_contact_sym_mig | 5 | -647.9 | 1305.8 | **-647.16** | 1304.32 | 482.33 | 0.00 | 203.4 | 6.197 | 2.745 | $\nu_1^a$ | $\nu_2^a$ | - | - | 0.037 | $m_{12}^b$ | 3.514 | 0.286 |
| sym_mig | 4 | -669.9 | 1347.8 | -669.95 | 1347.90 | 525.91 | 0.00 | 156.2 | 8.036 | 3.509 | $\nu_1^a$ | $\nu_2^a$ | 0.012 | $m_{12}^a$ | - | - | 5.306 | - |
| asym_mig | 5 | -677.0 | 1364.0 | **-669.27** | 1348.54 | 526.55 | 0.00 | 150.9 | 8.321 | 3.609 | $\nu_1^a$ | $\nu_2^a$ | 0.011 | 0.014 | - | - | 5.543 | - |
| anc_sym_mig | 5 | -671.7 | 1353.4 | **-670.04** | 1350.08 | 528.09 | 0.00 | 137.1 | 9.111 | 4.000 | $\nu_1^a$ | $\nu_2^a$ | 0.011 | $m_{12}^a$ | - | - | 6.202 | 0.000 |
| no_mig | 3 | -788.5 | 1583.0 | **-788.04** | 1582.08 | 760.09 | 0.00 | 345.1 | 3.911 | 1.607 | $\nu_1^a$ | $\nu_2^a$ | - | - | - | - | 1.716 | - |
| anc_asym_mig | 6 | -788.5 | 1589.0 | **-788.07** | 1588.14 | 766.15 | 0.00 | 340.5 | 3.968 | 1.635 | $\nu_1^a$ | $\nu_2^a$ | 0.000 | 0.000 | - | - | 0.052 | 1.692 |

Previous values of log-likelihood and AIC scores are from [2].
N — number of parameters.
$\log LL$ — log-likelihood.
* Inferred with *moments* with higher upper bounds of parameters.

Table S3: Demographic models for CVLN, CVLS populations of Gaboon forest frog *Scotobleps gabonicus*. Expected likelihood was received by using of ∂a∂i with $G = [40, 50, 60]$ grid size.

| | N | Prev. $\log LL$ | Prev. $AIC$ | $\log LL$ | $AIC$ | $\Delta AIC$ | $\omega_i$ | $\theta$ | $\nu_1^a$ | $\nu_2^a$ | $\nu_1^b$ | $\nu_2^b$ | $m_{12}^a$ | $m_{21}^a$ | $m_{12}^b$ | $m_{21}^b$ | $T_a$ | $T_b$ |
|---|---|---|---|---|---|---|---|---|---|---|---|---|---|---|---|---|---|---|
| unidir_asym_mig_size | 9 | - | - | -453.65 | 925.30 | 0.00 | 0.58 | 145.5 | 1.893 | 0.880 | 12.266 | 3.349 | - | 0.399 | 0.046 | 0.395 | 4.148 | 1.808 |
| sec_contact_asym_mig_size | 8 | -463.3 | 942.6 | **-455.17** | 926.34 | 1.04 | 0.34 | 240.8 | 1.072 | 0.679 | 7.383 | 1.948 | - | - | 0.071 | 0.722 | 1.207 | 1.139 |
| Structure 1,2 | 11 | - | - | -453.67 | 929.34 | 4.04 | 0.08 | 134.2 | 2.003 | 0.845 | 13.218 | 3.621 | 0.000 | 0.400 | 0.042 | 0.365 | 5.000 | 1.974 |
| unidir_sym_mig_size | 8 | - | - | -489.23 | 994.46 | 69.16 | 0.00 | 156.3 | 6.810 | 1.885 | 20.309 | 5.786 | - | 0.592 | 0.112 | $m_{12}^b$ | 3.129 | 0.313 |
| anc_asym_mig_size* | 8 | -519.6 | 1055.2 | **-499.16** | 1014.32 | 89.02 | 0.00 | 116.7 | 9.579 | 2.696 | 100 | 16.868 | 0.051 | 0.404 | - | - | 4.673 | 0.222 |
| anc_asym_mig_size | 8 | -519.6 | 1055.2 | **-500.47** | 1016.94 | - | - | 237.6 | 5.822 | 1.368 | 12 | 12 | 0.091 | 0.769 | - | - | 1.660 | 0.090 |
| sec_contact_asym_mig | 6 | -515.5 | 1043.0 | **-505.61** | 1023.22 | 97.92 | 0.00 | 255.2 | 6.209 | 1.738 | $\nu_1^a$ | $\nu_2^a$ | 0.106 | 0.707 | 0.950 | 0.530 | | |
| asym_mig | 5 | -513.5 | 1037.0 | **-512.96** | 1035.92 | 110.62 | 0.00 | 248.0 | 6.320 | 1.763 | $\nu_1^a$ | $\nu_2^a$ | 0.086 | 0.553 | - | - | 1.574 | - |
| anc_asym_mig | 6 | -520.8 | 1053.6 | **-512.98** | 1037.96 | 112.66 | 0.00 | 247.9 | 6.333 | 1.758 | $\nu_1^a$ | $\nu_2^a$ | 0.085 | 0.557 | - | - | 1.574 | 0.000 |
| sec_contact_sym_mig_size* | 7 | -537.9 | 1089.8 | **-513.12** | 1040.24 | 114.94 | 0.00 | 328.3 | 0.509 | 100 | 4.914 | 1.781 | - | - | 0.287 | $m_{12}^b$ | 0.301 | 0.777 |
| sec_contact_sym_mig_size | 7 | -537.9 | 1089.8 | **-514.07** | 1042.14 | - | - | 320.9 | 0.543 | 12 | 5.038 | 1.841 | - | - | 0.283 | $m_{12}^b$ | 0.319 | 0.799 |
| sec_contact_sym_mig | 5 | -553.8 | 1117.6 | **-551.78** | 1113.56 | 188.26 | 0.00 | 288.5 | 5.201 | 2.086 | $\nu_1^a$ | $\nu_2^a$ | - | - | 0.294 | $m_{12}^b$ | 0.767 | 0.398 |
| anc_sym_mig_size | 7 | -600.8 | 1215.6 | **-550.34** | 1114.68 | 189.38 | 0.00 | 254.8 | 4.440 | 2.179 | 12 | 2.810 | 0.323 | $m_{12}^a$ | - | - | 1.447 | 0.112 |
| sym_mig | 4 | -556.1 | 1120.2 | **-555.37** | 1118.74 | 193.44 | 0.00 | 268.9 | 5.459 | 2.165 | $\nu_1^a$ | $\nu_2^a$ | 0.228 | $m_{12}^a$ | - | - | 1.342 | - |
| anc_sym_mig_size* | 7 | -600.8 | 1215.6 | **-553.00** | 1120.00 | - | - | 72.7 | 10.319 | 7.245 | 54.547 | 9.098 | 0.143 | $m_{12}^a$ | - | - | 8.012 | 0.738 |
| anc_sym_mig | 5 | -558.3 | 1126.6 | **-555.38** | 1120.76 | 195.46 | 0.00 | 265.6 | 5.521 | 2.192 | $\nu_1^a$ | $\nu_2^a$ | 0.227 | $m_{12}^a$ | - | - | 1.368 | 0.000 |
| no_mig_size* | 5 | -704.6 | 1419.2 | **-691.47** | 1392.94 | 467.64 | 0.00 | 464.3 | 1.797 | 100 | 4.510 | 1.179 | - | - | - | - | 0.137 | 0.292 |
| no_mig_size | 5 | -704.6 | 1419.2 | **-692.26** | 1394.52 | 469.22 | 0.00 | 465.6 | 1.805 | 12 | 4.530 | 1.192 | - | - | - | - | 0.137 | 0.288 |
| no_mig | 3 | -704.4 | 1414.8 | -704.35 | 1414.70 | 489.40 | 0.00 | 463.7 | 4.050 | 1.407 | $\nu_1^a$ | $\nu_2^a$ | - | - | - | - | 0.411 | - |

Previous values of log-likelihood and AIC scores are from [2].

N — number of parameters.

$\log LL$ — log-likelihood.

* Inferred with *moments* with higher upper bounds of parameters.

Table S4: Demographic models for Northern, Southern populations of Gaboon forest frog *Scotobleps gabonicus*. Expected likelihood was received by using of ∂a∂i with $G = [50, 60, 70]$ grid size.

| | N | Prev. $\log LL$ | Prev. $AIC$ | $\log LL$ | $AIC$ | $\Delta AIC$ | $\omega_i$ | $\theta$ | $\nu_1^a$ | $\nu_2^a$ | $\nu_1^b$ | $\nu_2^b$ | $m_{12}^a$ | $m_{21}^a$ | $m_{12}^b$ | $m_{21}^b$ | $T_a$ | $T_b$ |
|---|---|---|---|---|---|---|---|---|---|---|---|---|---|---|---|---|---|---|
| unidir_asym_mig_size | 9 | - | - | -365.29 | 748.58 | 0.00 | 0.44 | 251.9 | 0.139 | 6.899 | 0.889 | 8.873 | 2.639 | - | 0.556 | 0.312 | 1.089 | 0.109 |
| unidir_sym_mig_size | 8 | - | - | -365.31 | 748.62 | 0.04 | 0.43 | 249.1 | 0.177 | 6.876 | 1.164 | 10.172 | 2.117 | - | 0.424 | $m_{12}^b$ | 1.135 | 0.085 |
| anc_asym_mig_size | 8 | -379.8 | 775.6 | **-368.22** | 752.44 | 3.86 | 0.06 | 248.3 | 0.224 | 7.034 | 12 | 12 | 1.849 | 0.158 | - | - | 1.184 | 0.044 |
| anc_asym_mig_size* | 8 | -379.8 | 775.6 | **-368.23** | 752.46 | - | - | 240.1 | 0.271 | 6.826 | 100 | 54.277 | 1.487 | 0.154 | - | - | 1.263 | 0.039 |
| Structure 1,2 | 11 | - | - | -365.26 | 752.52 | 3.94 | 0.06 | 250.6 | 0.149 | 7.034 | 0.974 | 8.707 | 2.510 | 0.001 | 0.507 | 0.328 | 1.110 | 0.101 |
| sec_contact_asym_mig_size | 8 | -379.4 | 774.8 | **-369.84** | 755.68 | 7.10 | 0.01 | 259.7 | 0.010 | 6.522 | 0.436 | 7.497 | - | - | 1.206 | 0.169 | 0.729 | 0.383 |
| sec_contact_asym_mig | 6 | -378.0 | 768.0 | **-374.56** | 761.12 | 12.54 | 0.00 | 264.7 | 0.369 | 7.076 | $\nu_1^a$ | $\nu_2^a$ | - | - | 1.325 | 0.226 | 0.752 | 0.305 |
| asym_mig | 5 | -379.1 | 768.2 | **-377.82** | 765.64 | 17.06 | 0.00 | 256.0 | 0.390 | 7.172 | $\nu_1^a$ | $\nu_2^a$ | 1.014 | 0.168 | - | - | 1.145 | - |
| anc_asym_mig | 6 | -379.8 | 771.6 | **-377.83** | 767.66 | 19.08 | 0.00 | 259.1 | 0.382 | 7.110 | $\nu_1^a$ | $\nu_2^a$ | 1.038 | 0.168 | - | - | 1.121 | 0.000 |
| sec_contact_sym_mig_size* | 7 | -412.4 | 838.8 | **-399.99** | 813.98 | 65.40 | 0.00 | 260.3 | 100 | 5.267 | 0.543 | 6.949 | - | - | 0.364 | $m_{12}^b$ | 0.508 | 0.597 |
| sec_contact_sym_mig_size | 7 | -412.4 | 838.8 | **-400.10** | 814.20 | - | - | 261.7 | 12 | 5.539 | 0.531 | 6.936 | - | - | 0.372 | $m_{12}^b$ | 0.525 | 0.565 |
| sec_contact_sym_mig | 5 | -406.4 | 822.8 | **-405.35** | 820.70 | 72.12 | 0.00 | 305.2 | 0.576 | 6.313 | $\nu_1^a$ | $\nu_2^a$ | - | - | 0.739 | $m_{12}^b$ | 0.645 | 0.122 |
| sym_mig | 4 | -410.5 | 829.0 | -410.45 | 828.90 | 80.32 | 0.00 | 266.2 | 0.638 | 6.672 | $\nu_1^a$ | $\nu_2^a$ | 0.355 | $m_{12}^a$ | - | - | 1.071 | - |
| anc_sym_mig | 5 | -411.6 | 833.2 | **-410.44** | 830.88 | 82.30 | 0.00 | 267.0 | 0.635 | 6.659 | $\nu_1^a$ | $\nu_2^a$ | 0.356 | $m_{12}^a$ | - | - | 1.065 | 0.000 |
| anc_sym_mig_size | 7 | -411.3 | 836.6 | **-410.44** | 834.88 | 86.30 | 0.00 | 268.2 | 0.621 | 6.593 | 12 | 12 | 0.365 | $m_{12}^a$ | - | - | 1.061 | 0.002 |
| anc_sym_mig_size* | 7 | **-411.3** | 836.6 | -443.84 | 901.68 | - | - | 67.8 | 4.208 | 18.898 | 1.723 | 35.187 | 1.346 | $m_{12}^a$ | - | - | 7.217 | 0.995 |
| no_mig_size* | 5 | -533.9 | 1077.8 | **-531.74** | 1073.48 | 324.90 | 0.00 | 411.0 | 100 | 100 | 0.312 | 4.568 | - | - | - | - | 0.165 | 0.194 |
| no_mig_size | 5 | **-533.9** | 1077.8 | -535.43 | 1080.86 | - | - | 414.7 | 12 | 12 | 0.313 | 5.061 | - | - | - | - | 0.164 | 0.191 |
| no_mig | 3 | -549.1 | 1104.2 | -549.12 | 1104.24 | 355.66 | 0.00 | 439.3 | 0.432 | 5.904 | $\nu_1^a$ | $\nu_2^a$ | - | - | - | - | 0.292 | - |

Previous values of log-likelihood and AIC scores are from [2].

N — number of parameters.

$\log LL$ — log-likelihood.

* Inferred with *moments* with higher upper bounds of parameters.
