## Supplementary material for "GADMA: Genetic algorithm for inferring demographic history of multiple populations from allele frequency spectrum data": All data, parameters for GADMA runs and results

All results, AFS data, scripts and parameters for GADMA that were used in the paper is openly accessible via the link: `https://bitbucket.org/noscode/gadma_results`.
